# Impaired HPV driven CD8 T cells recognition and Immune suppression in HPV-Induced Cervical Cancer

**DOI:** 10.64898/2026.01.19.700360

**Authors:** Mohammad Kadivar, Dorthe Blirup Snejbjerg, Marie C Viuff, Grigori Nos, Stine Kiær Larsen, Jesper Bonde, Benny Kirschner, Susanne K. Kjær, Kirsten Marie Jochumsen, Sine R Hadrup

## Abstract

Human papillomavirus (HPV) remains the leading cause of cervical cancer, yet the mechanisms underlying its immune evasion remain poorly defined. We performed an integrated analysis of HPV-specific CD8 T cell responses and the immune microenvironment in cervical cancer, high-grade intraepithelial neoplasia (CIN3), and healthy controls using flow cytometry, transcriptomics, and DNA-barcoded peptide–MHC multimer screening across cervical biopsies, peripheral blood, and liquid-based cytology (LBC).

Cervical cancer tissues exhibited a profoundly immunosuppressive milieu, with enrichment of exhausted CD8 and CD4 T cells, increased regulatory T cells, and PD-L1–expressing myeloid subsets. Conventional dendritic cells and macrophages showed reduced frequencies, while plasmacytoid dendritic cells and intermediate monocytes were elevated. Interestingly, LBC samples reliably reflected T cell exhaustion signatures observed in biopsies, supporting their use as a minimally invasive tool for T cell immune monitoring; however, they were not suitable for detailed myeloid profiling. Transcriptomic analysis revealed distinct gene expression profiles in tumor tissues with elevated signatures of immune checkpoints and regulatory immune cell infiltration.

Importantly, HPV-specific CD8 T cell responses were significantly reduced in cancer patients compared to CIN3 and controls, with decreased breadth and frequency of HPV peptides recognition. HPV peptides screening identified HPV-specific CD8 T cell responses toward 109 unique peptide-MHC complexes, including 37 novel HPV-derived peptide from E2, E6, and E7 proteins.

These findings reveal impaired HPV immune recognition and a suppressive tumor microenvironment in cervical cancer, underscoring the need to enhance HPV-specific T cell responses and target immune suppression in therapeutic strategies.

## Introduction

Despite the availability of an efficacious prophylactic human papillomavirus (HPV) vaccine there is still a considerable global burden of HPV-related diseases. HPV is the most common viral infection of the female reproductive tract with preference for epithelial cells and is the primary cause of cervical cancer.^1^ In addition to cervical cancer, HPV infection is implicated in several other cancer types, including head and neck cancers. Cervical cancer is the fourth most common malignancy diagnosed in women worldwide, accounting for 3.1% of new cancer cases and 3.3% of cancer-related deaths across all ages and both sexes, according to WHO data from 2020.^2–4^ It is estimated that more than 80% of women are infected with HPV during their lifetime. While the majority of HPV infections are cleared by the immune system within 6–12 months, a small proportion remains as persistent infections,^5^ which can lead to the development of cervical intraepithelial neoplasia and cancer. Only about 3–5% of cervical HPV infections progress to transforming infections. Among patients with high-grade cervical intraepithelial neoplasia (CIN3), the cumulative risk of developing cervical cancer is approximately 30% over a 30-year period if left untreated. With appropriate treatment, this risk drops to below 1%, although it still remains higher than that observed in the general female population.^6–8^ Particularly, HPV-16 and -18 are considered having the highest oncogenic potential and are reported to account for approximately >70% of cervical cancer cases.^9,10^

The HPV genome is a circular double-stranded DNA molecule that encodes six early genes (E1, E2, E4, E5, E6 and E7) and two late genes (L1 and L2), along with a non-coding region. Among these, the E6 and E7 genes are particularly significant due to their roles in inactivating host tumour suppressor genes. Continuous expression of E6 and E7 in HPV-16 and -18 is required to induce and maintain the neoplastic phenotype and drive oncogenic progression.^5,11,12^ The E2 gene functions as a transcriptional repressor of E6 and E7; however, when the viral DNA integrates into the host genome, the E2 sequence is often disrupted, resulting to increased expression of E6 and E7. E2 therefore plays a critical role in oncogenic progression of HPV.^5,13,14^ Based on these roles, E2, E6, and E7 were selected to investigate the epitope-specific recognition of these HPV proteins by CD8 T cells in this study.

The immune system plays a key role in controlling HPV infection, with cytotoxic CD8 T cells being particularly crucial for clearing infected and cancerous cells.^15^ However, high-risk HPV types employ complex immune evasion mechanisms during the development of cervical cancer. Understanding these mechanisms is essential for the development of targeted therapies and immunotherapeutic approaches for HPV-associated malignancies.

This study aimed to comprehensively map HPV-specific CD8 T cells recognizing HPV types 16 and 18; and to characterize differences in T cell recognition profiles between healthy controls and patients with CIN3 or cervical cancer. Furthermore, the study aimed to explore alterations in both local and systemic immune infiltration and the microenvironment in CIN3 and cervical cancer, focusing on phenotypic markers of CD8 and CD4 T cells, including activation, differentiation, and exhaustion states. Additionally, innate immune responses and changes in the myeloid cell compartment were investigated to identify cellular signatures and phenotypic shifts associated with oncogenic transformation and cancer progression.

## Results

Samples were collected from three distinct groups of individuals: healthy controls (i.e. women without cervical neoplasia), patients diagnosed with CIN3, and patients with cervical cancer. For each participant, three different types of samples were obtained: peripheral blood, cervical biopsies, and cervical liquid-based cytology (LBC) samples (Figure 1).

**Figure 1:**
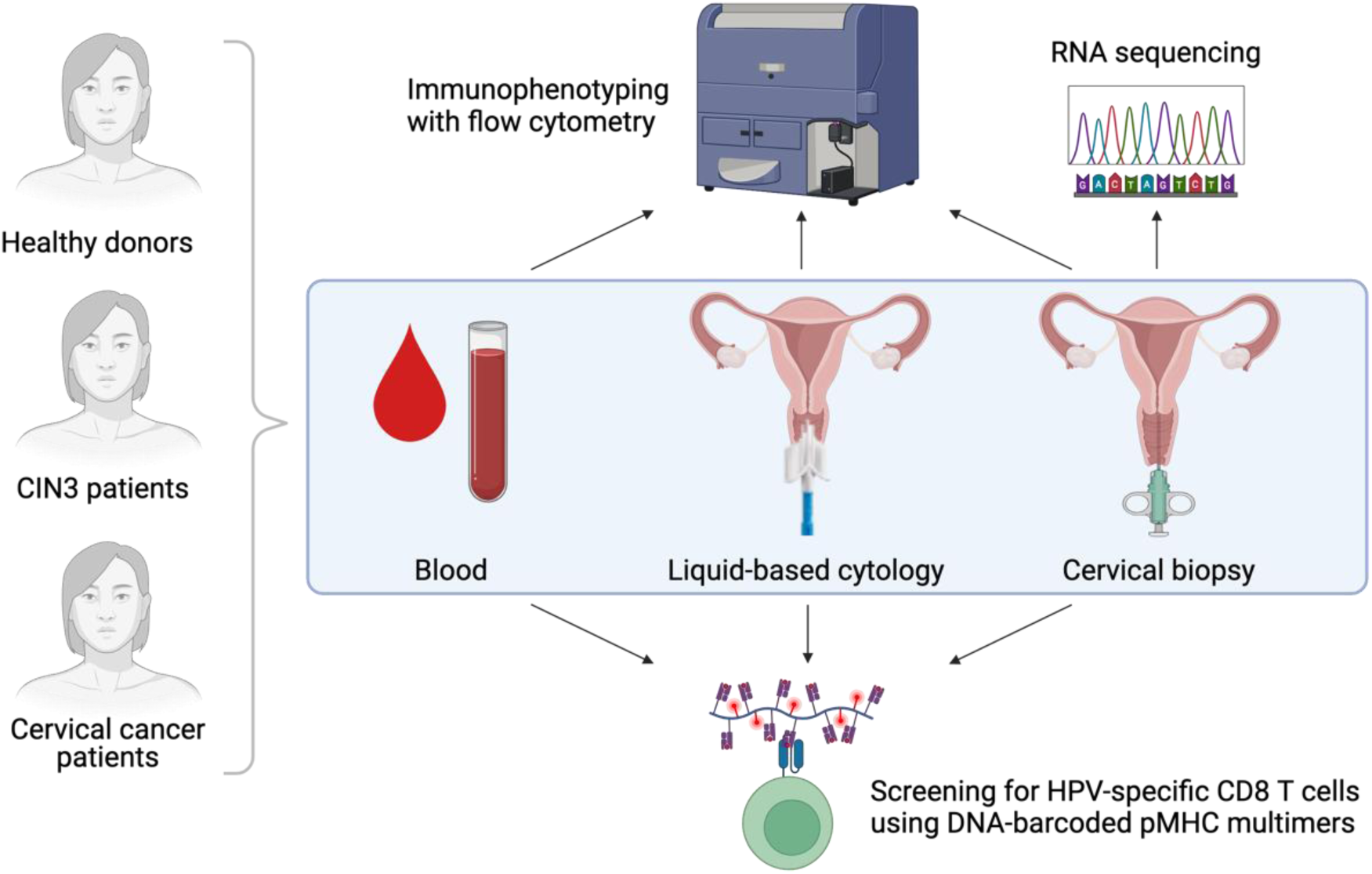
Experimental Design and Sample Collection. Blood, cervical biopsies; and liquid-based cytology (LBC) samples were obtained from healthy control donors and patients with CIN3 or cervical cancer. These samples were analyzed by flow cytometry for immunophenotyping and/or screened using a HPV derived peptides library to detect HPV-specific CD8 T cell responses. In addition, RNA sequencing was performed on cervical biopsy samples.

### Tumor infiltrating CD8 T cells showed exhaustion signature

To characterize tumor infiltrating CD8 T cells and compare them with tissue infiltrating CD8 T cells in CIN3 and control groups, we performed high-dimensional clustering analysis on our multi-color flow cytometry data. Cervical biopsy samples from ten individuals in each group were analyzed and visualized using UMAP dimensionality reduction (Figure 2A). FlowSOM clustering analysis of CD8 T cells from these biopsies identified eight distinct clusters (Supplementary Figure 2A). The percentage of CD39^+^CD103^+^PD-1^+^CD27^+^ CD8 T_EM_ clusters (FlowSOM clusters Pop6 and Pop7) was higher in the cervical biopsies from cancer group compared to the CIN3 and control groups. Both CD8 T_EM_ clusters expressed TOX and intermediate level of EOMES. Similar to cervical biopsies, the percentage of CD39^+^CD103^+^PD-1^+^CD27^+^ CD8 T_EM_ clusters (FlowSOM clusters Pop1 and Pop4) was higher in the LBC samples from the cancer group compared to the CIN3 and control groups (Figure 2B). Although CD39^+^CD103^+^ CD8 T cells are known to display an exhausted and tissue-resident memory (T_RM_) gene signature,^16,17^ these cells also express Ki67, indicating that they are in a proliferating state (Supplementary Figure 2A). The two CD8 T_EM_ clusters in cervical biopsies differed only in CD57 expression, with Pop7 being CD57^+^CD39^+^CD103^+^PD-1^+^CD27^+^ CD8 T_EM_, while Pop6 cluster had no expression of CD57. Interestingly, only the percentage of CD57^+^CD39^+^CD103^+^PD-1^+^CD27^+^ CD8 T_EM_ cells correlated with cancer stage (Supplementary Figure 3C).

**Figure 2:**
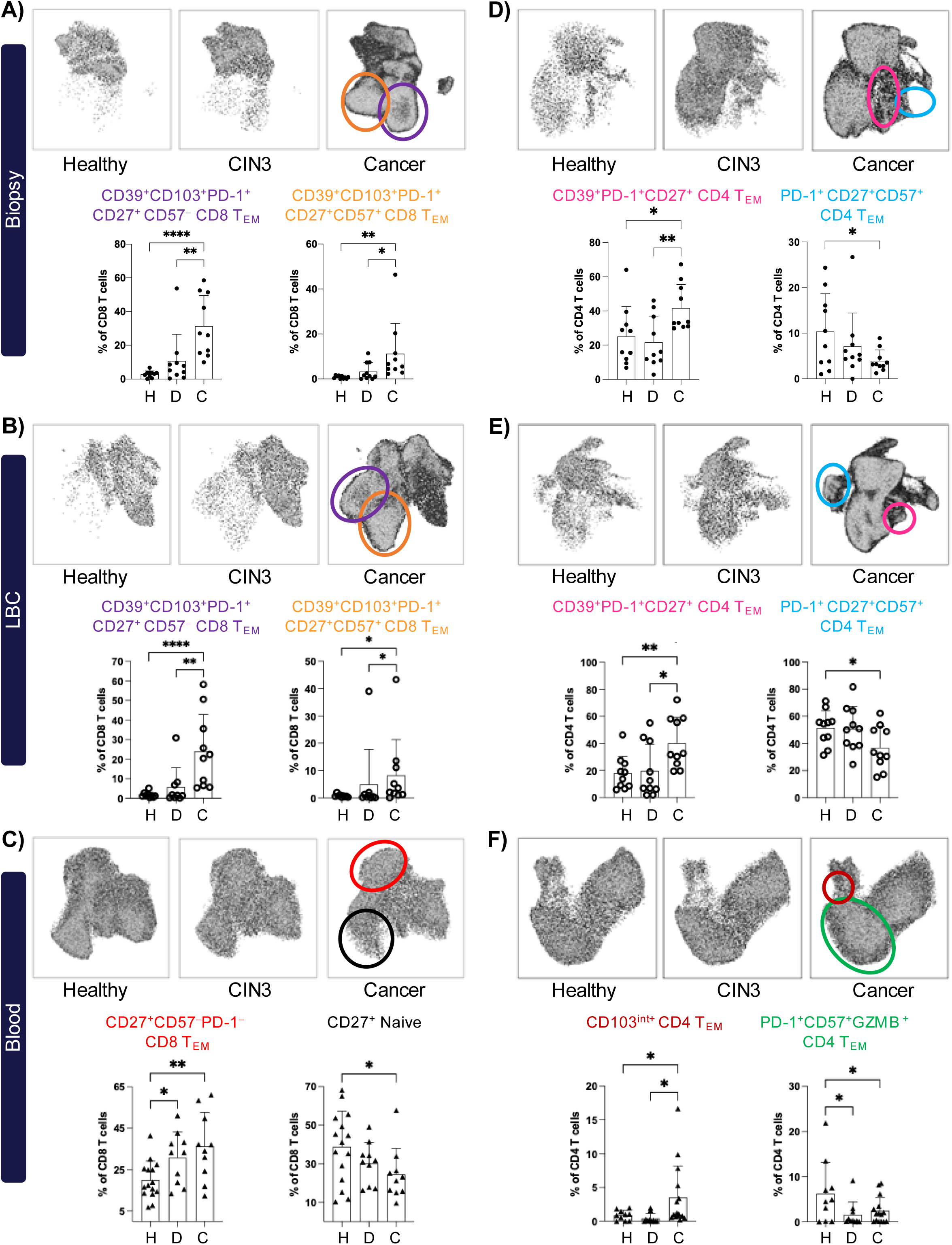
Presence of exhaustion signature in tumor infiltrating T Cells. UMAP illustration of both CD8 (A-C) and CD4 (D-F) T cells in control, CIN3, and cancer groups. T cells were clustered into 8 different subpopulations using FlowSOM. Filled circles represent values from cervical biopsy samples, open circles from liquid-based cytology (LBC), and triangles from blood. A) In the cervical biopsies, the frequency of CD39^+^PD-1^+^CD103^+^CD27^+^ CD8 T_EM_ cell clusters were significantly higher in patients with cervical cancer compared to CIN3 and control groups. B) In the LBC samples, comparable to biopsies; the frequency of CD39^+^PD-1^+^CD103^+^CD27^+^ CD8 T_EM_ cell clusters were higher in patients with cancer compared to CIN3 and control groups. C) In the blood, the frequency of CD27^+^CCR7^−^CD45RA^−^ CD8 T cell cluster were higher in patients with cancer compared to CIN3 and control groups. The frequency of CD27^+^CCR7^+^CD45RA^+^ CD8 T cell cluster were lower in patients with cancer compared to CIN3 and control groups. D) In the cervical biopsies, the frequency of CD39^+^PD-1^+^CD27^+^CD103^−^ CCR7^−^CD45RA^−^ CD4 T cell cluster was significantly higher in patients with cervical cancer compared to CIN3 and control groups. However, the frequency of CD39^−^PD-1^+^CD103^−^CD27^+^CD57^+^CCR7^−^CD45RA^−^ CD4 T cell cluster was lower in patients with cervical cancer compared to CIN3 and control groups. E) In the LBC samples, the frequency of CD39^+^PD-1^+^CD27^+^CD103^−^CCR7^−^CD45RA^−^ CD4 T cell cluster was higher in patients with cancer compared to CIN3 and control groups. The frequency of CD39^−^PD-1^+^CD103^−^CD27^+^CD57^−^CCR7^−^CD45RA^−^ CD4 T cell cluster was lower in patients with cervical cancer compared to CIN3 and control groups. F) In the blood, the frequency of CD103^int^ CCR7^−^CD45RA^−^ CD4 T cell cluster was higher in patients with cancer compared to CIN3 and control groups. The frequency of CD57^+^GZMB^+^PD-1^+^CCR7^−^CD45RA^−^ CD4 T cell cluster was higher in control group compared to CIN3 and cancer groups. Plots show mean ± SD (Kruskal-Wallis uncorrected Dunn’s test, *: p<0.05, **: p<0.01, ***: p< 0.001).

Although most CD8 T cells in the cervical biopsies were CCR7^−^CD45RA^−^ CD8 T cells (CD8 T_EM_), the percentage of total CD8 T_EM_ cells was similar in the biopsies of all groups (Supplementary Figure 3A). The percentage of single positive CD39, EOMES, or TOX CD8 T cells were higher in cancer patients compared to CIN3 and controls showing an exhausted signature of CD8 T cells in the cervical cancer biopsies compared to non-cancer cervical tissues (Supplementary Figure 3B). The percentage of Ki67^+^ CD8 T cells was also higher in the cancer biopsies compared to CIN3 and normal biopsies (Supplementary Figure 3B).

### Cervical cancer patients have higher percentage of circulating CD27^+^ CD8 T_EM_ cells

Furthermore, to characterize the circulating CD8 T cells in the different groups, we performed high-dimensional clustering analysis on the flow cytometry data derived from analyses for the peripheral blood. Blood samples from thirty-five individuals were examined in total and visualized using UMAP dimensionality reduction (Figure 2C). By using FlowSOM clustering analysis on the CD8 T cells from blood samples, eight different clusters were identified (Figure 2C). The CD8 T_EM_ cells (CCR7^−^CD45RA^−^ CD8 T cell) were divided into 3 clusters: Pop0, Pop1, and Pop7. However, only the percentage of CD27^+^PD-1^−^CD57^−^GZMB^−^Ki67^−^ CD8 T_EM_ subset (Pop1) was higher in the cancer group compared to that in CIN3 and in controls (Figure 2C). The percentage of more differentiated T_EM_ subset (CD27^−^PD-1^−^CD57^+^GZMB^+^Ki67^−^ CD8 T_EM_ subset – Pop7), and the proliferating CD8 T_EM_ subset (CD27^+^PD-1^+^CD57^−^GZMB^−^Ki67^+^ CD8 T_EM_ subset – Pop0), were similar in the three study groups (Figure 2C). We also observed a trend towards the percentage of single positive EOMES or TOX CD8 T cells were lower in cancer patients compared to in patients with CIN3 and controls, showing the less differentiated signature of CD8 T cells in the cancer patients compared to the CIN3 and control groups (Supplementary Figure 2B).

Moreover, the frequency of naïve CD8 T cells (CD27^+^CCR7^+^CD45RA^+^ CD8 T cell cluster – Pop2) were lower in patients with cancer compared to the groups of CIN3 or controls (Figure 2C).

### Presence of exhaustion signature in tumor infiltrating CD4 T Cells

We performed the same analysis on tissue-infiltrating CD4 T cells as we did on CD8 T cells to investigate the changes in CD4 T cells among the controls, CIN3 patients, and cancer patients. The cervical biopsy samples from ten individuals were examined from each group and visualized using UMAP dimension reduction (Figure 2D). By using FlowSOM clustering analysis on the CD4 T cells from biopsies, eight different clusters were identified (Supplementary Figure 2D). Out of all CD39^+^ CD4 T_EM_ cells, the percentage of CD39^+^PD-1^+^CD27^+^ CD4 T_EM_ cells (cluster Pop5) was significantly higher in the cancer group compared to in the CIN3 and control groups (Figure 2D). In the LBC samples, similar to biopsy samples, the frequency of CD39+PD-1+CD27+CD103− CD4 T_EM_ cluster was higher in patients with cancer compared to CIN3 and controls (Figure 2E). The percentage of CD39−PD-1+CD103−CD27+CD57+ CD4 T_EM_ cell cluster was lower in patients with cervical cancer compared to CIN3 and control groups (Figure 2D). In contrast with CD8 T cells, the triple positive (CD39+CD103+PD-1+) CD4 TEM cells were not enriched in the tumour biopsies compared to non-tumour biopsies. However, the percentage of single positive CD39 or EOMES CD4 T cells were higher in cancer patients compared to CIN3 and controls showing the exhausted signature of CD4 T cells in the cervical tumour biopsies compared to non-tumour cervical tissues (Supplementary Figure 3B). The percentage of TCF-1^+^ CD4 T cells was also higher in the cancer biopsies compared to CIN3 and normal biopsies (Supplementary Figure 3B). Similar to CD8 T cells, most of CD4 T cells in the cervical biopsies are CCR7^−^CD45RA^−^ effector memory CD8 T cells (CD8 T_EM_), and the percentage of total CD4 T_EM_ cells was similar in the biopsies of all groups (Supplementary Figure 3A).

In the blood, the percentage of PD-1+CD57+GZMB + CD4 T_EM_ cell cluster was lower in CIN3 and cancer groups compared to the control group (Figure 2F). Moreover, the percentage of CD103^int^ CD27^−^ CD4 T_EM_ cell cluster was higher in patients with cancer compared to CIN3 and control groups (Figure 2F). The percentage of single positive EOMES CD4 T cells were lower in cancer patients compared to CIN3 and controls (Supplementary Figure 3B).

Overall, the flow cytometry data on T cells revealed that both CD8 and CD4 T cells in cervical cancer patients exhibit higher levels of exhaustion markers compared to those in CIN3 and controls. This suggests that T cell exhaustion is a significant feature of the immune landscape in cervical cancer, potentially contributing to the tumor’s ability to evade immune surveillance.

Our study also importantly found a consistent pattern of T cell exhaustion markers in both cervical LBC samples and cervical biopsies. This similarity indicates that LBC samples can reliably reflect the T cell immune profile observed in biopsies. The major advantage of this finding is that LBC sampling is less invasive than biopsies, making it a more patient-friendly option for monitoring T cell responses and potentially diagnosing cervical cancer.

### Altered innate immune response in blood and cervical samples of cancer patients

In our investigation of the immune cell profiles in cancer patients, we observed significant differences in the distribution of circulating conventional dendritic cells (cDCs) compared to controls. The percentage of cDCs in the peripheral blood of cancer patients was lower than in patients with CIN3 or controls (Figure 3A). A similar trend was noted in biopsy samples from tumor tissues, although the difference did not reach statistical significance (Figure 3B). This reduction in cDCs in both blood and tissue suggests a compromised ability to present antigen, thereby impairing the initiation of effective anti-tumor immune responses in cancer patients.

**Figure 3:**
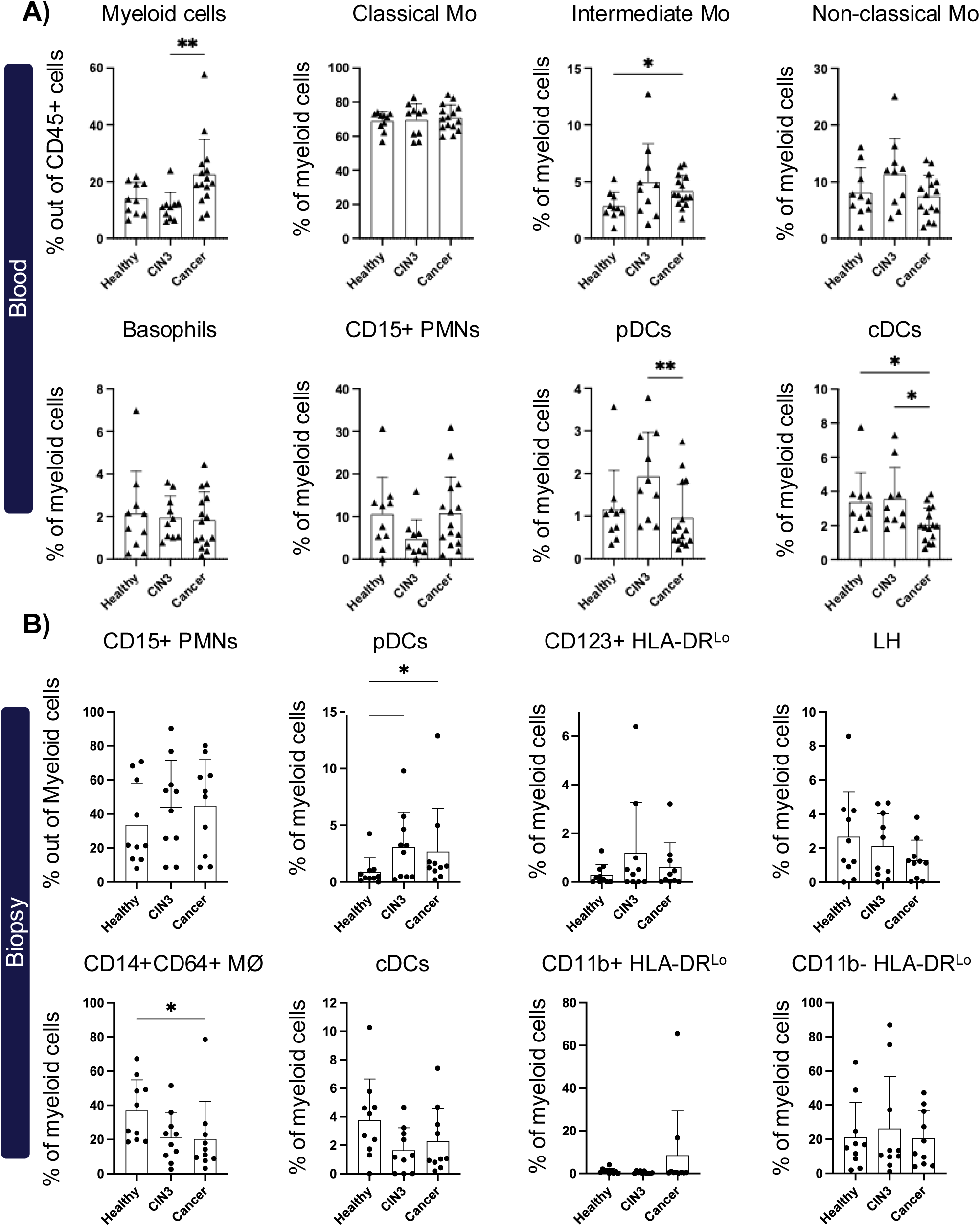
Altered innate immune response in blood and cervical samples of cancer patients. The percentage of different myeloid cells in biopsy (A) and blood (B) of healthy control donors and patients with CIN3 or cervical cancer. The myeloid cell compartment in tissue differs from that in blood, so we used distinct definitions for each myeloid cell subset to investigate their frequencies. Plots show mean ± SD (Kruskal-Wallis uncorrected Dunn’s test, *: p<0.05, **: p<0.01, ***: p< 0.001). Filled circles represent values from cervical biopsy samples and triangles from blood.

On the other hand, plasmacytoid dendritic cells (pDCs) exhibited a higher percentage in the cervix of both cancer patients and those with CIN3 compared to controls (Figure 3B). The increased presence of pDCs, which are known for their role in antiviral responses and induction of immune tolerance, may contribute both to the anti-HPV immune response and to the development of an immunosuppressive microenvironment. This dual role could potentially facilitate tumor progression and persistence.

Furthermore, our analysis of cervical tissue samples revealed that CD14^+^CD64^+^ macrophages were significantly less frequent in cancer tissues compared to controls (Figure 3B). However, in the peripheral blood, only the percentage of intermediate monocytes were higher in cancer patients compared to controls (Figure 3A). The other subsets of myeloid cells did not show any differences between different groups, however, the frequency of myeloid cells out of all hematopoetic cells (CD45^+^ cells) was higher in the peripheral blood of cancer patients compared to other groups (Figure 3A).

Altogether, in the myeloid compartment, we observed a reduction in cDCs and CD14^+^CD64^+^ macrophages, which may impair effective immune responses. Additionally, there was an increase in pDCs and intermediate monocytes, potentially contributing to an immunosuppressive environment.

In cervical LBC samples, the myeloid compartment was underrepresented compared with cervical biopsies. Several subsets were detected only at very low frequencies, including pDCs, Langerhans (LH) cells, CD123⁺HLA-DR^low^ cells, and CD11b⁻HLA-DR^low^ cells (Supplementary Figure 5). The overall percentage of other myeloid populations was also lower than in biopsy samples (Supplementary Figure 5). Despite the overall differences in myeloid representation between sample types, both LBC samples and cervical biopsies demonstrated a comparable pattern, with cDC frequencies consistently decreased in the cancer group relative to the control group (Supplementary Figure 5). These observations suggest that, while LBC samples reflect some aspects of the cervical myeloid landscape, biopsy specimens provide a more comprehensive representation of myeloid populations and therefore remain the preferred sample type for detailed myeloid profiling.

### Immunosuppressive microenvironment in the cervix of patients with cancer

To evaluate the presence of an immunosuppressive tumor microenvironment in the cancer patients, we examined the frequency of regulatory T cells and the level of PDL-1 expression by myeloid cells in the cervical samples from all the three groups. Our results demonstrated a significant increase in the frequency of regulatory T cells (CD4^+^CD25^+^CD127^−^ Tregs) within tumor tissues compared to CIN3 and control cervical samples (Figure 4A). This was accompanied by higher expression of PD-L1 on various myeloid cell subsets (including CD14^+^CD64^+^ macrophages, cDCs, pDCs, and CD11b^+^HLA-DR^lo^ cells) in the tumor microenvironment compared to CIN3 and control cervical tissues (Figure 4B). Moreover, we observed that the expression of CD103 is higher on cDCs, CD11b^−^HLA-DR^lo^, and CD14^+^CD64^+^ macrophages in tumor samples compared to cervical tissues of CIN3 and controls (Supplementary Figure 5A). Interestingly, when analyzing the peripheral blood samples, we found that the frequency of Tregs was similar between cancer patients, CIN3, and controls (Figure 4A). These findings suggest that the tumor microenvironment specifically fosters a more immunosuppressive milieu, characterized by increased Tregs and elevated PD-L1 and CD103 expression on myeloid cells, which could contribute to immune evasion by the tumour. Conversely, the systemic immune profile, as reflected in peripheral blood, does not exhibit these alterations, highlighting the localized nature of immune modulation within the tumour site.

**Figure 4:**
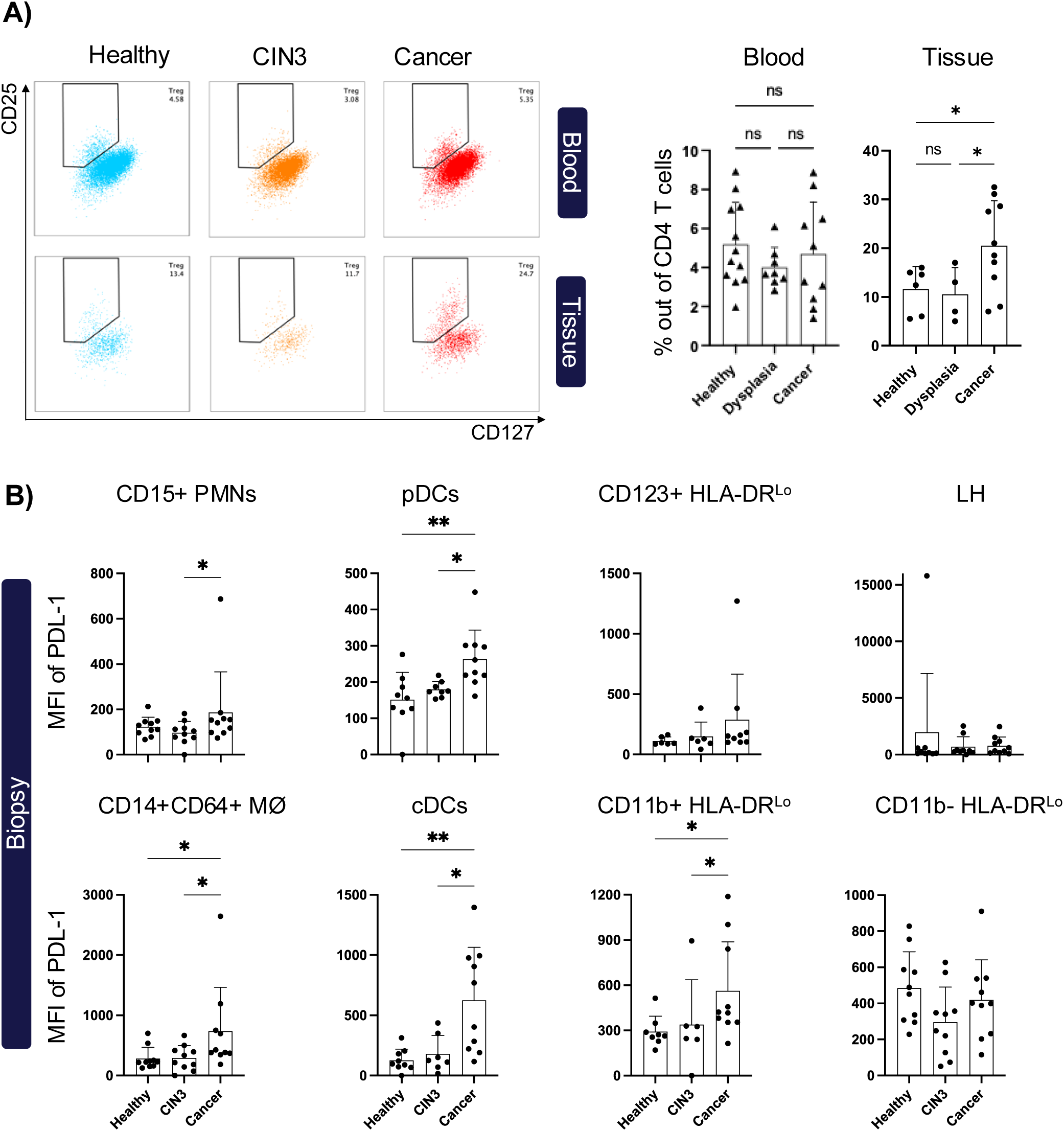
Presence of immunosuppressive microenvironment in the cervix of patients with cancer. A) Tregs are enriched in the tissue of cancer patients compared to other groups. CD25+CD127− CD4 T cells were defined as Tregs. B) The expression (MFI) of PDL-1on different subsets of cervical myeloid cells from biopsy samples is shown. Plots show mean ± SD (Kruskal-Wallis uncorrected Dunn’s test, *: p<0.05, **: p<0.01, ***: p< 0.001). MFI: mean fluorescence intensity. Filled circles represent values from cervical tissue samples and triangles from blood.

### Assessment of CD8 T cell recognition to HPV- E2, E6, E7 derived HLA ligands

To investigate the T cell response to HPV, we examined CD8 T cell recognition towards MHC-I binding peptides from the oncogene proteins E6 and E7 along with the E2 protein gene of both HPV16 and HPV18. Using NetMHCpan 4.0, we selected 685 distinct MHC-binding peptides (8 to 11 amino acids) in our peptide library for screening. These peptides were predicted to bind to one or more of fourteen prevalent MHC-I molecules, including HLA-A (A01:01, A02:01, A03:01, A11:01, A24:02), HLA-B (B07:02, B08:01, B15:01, B35:01) and HLA-C (C03:04, C04:01, C05:01, C07:01, C07:02) loci; leading to a total >1100 peptide-MHC specificities for experimental evaluation (Figure 5A). Given that the E2 protein is larger compared to the E6 and E7 proteins, it accounted for approximately 60% of the total predicted peptides. In this study, CD8 T cell reactivity to these peptides was evaluated using the three different sample types: PBMCs from 39 subjects, cervical biopsies from 16 subjects, and LBC specimens from 17 subjects (Supplementary Table 1). Patients were screened for CD8 T cell reactivity according to their HLA-type. We covered 14 frequent HLA haplotypes in the screen, providing a mean HLA coverage of 3.3 HLA per patient, and an average of 276 pMHC multimers were used per patient. To screen for CD8 T cell reactivity against HPV, briefly, each pMHC complex was multimerized on a dextran backbone labelled with Phycoerythrin (PE) or Allophycocyanin (APC) and tagged with a unique DNA barcode. These DNA-barcoded pMHC multimers were pooled to create a patient-specific pMHC multimer panel tailored to the individual’s HLA profile. This panel was then incubated with T cells derived from the biospecimens from the patients. CD8 T cells binding pMHC multimers were subsequently sorted and sequenced to identify T cell recognition within such samples. We also included 39 T cell epitopes from common viruses, specifically cytomegalovirus (CMV), Epstein-Barr virus (EBV), and influenza (Flu) virus, collectively referred to as the CEF epitopes (Figure 5A).

**Figure 5:**
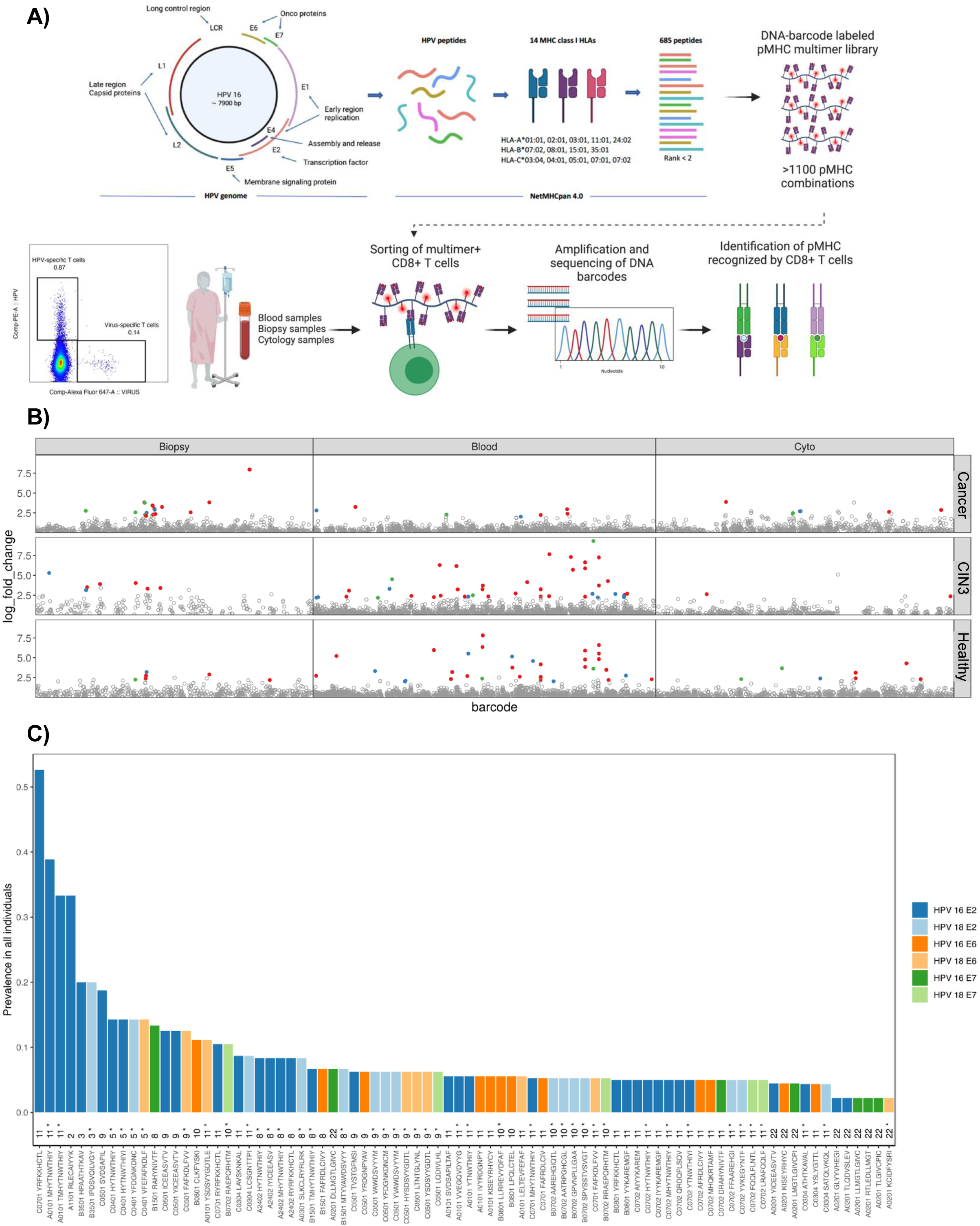
Mapping of HPV CD8 T cell epitopes. A) Schematic representation of the complete HPV 16 circular genome with all genes represented. The early region E2, E6 and E7 genes of HPV 16/18 are used for identification of 685 peptides with predicted binding rank of <2 for 14 prevalent HLA-A, HLA-B and HLA-C alleles using NetMHCpan 4.0. The total of 1238 unique peptide-MHC (pMHC) pairs were constructed for making peptide libraries to screen for HPV-specific CD8 T cells. T cells then stained with pMHC multimers and multimer+ CD8 T cells that were sorted for DNA barcode sequencing analysis to identify epitope recognition. Representative flow cytometry plot of pMHC multimer staining of CD8 T cells from an HPV positive patient showing PE+ (HPV-specific) and APC+ (CEF-specific) CD8 T cells. B) Dot-plots showing summary of all T cell recognition to HPV derived peptides identified in the blood, biopsy, and LBC of individuals in control, CIN3, and cancer groups. CD8 T cell recognition to individual epitopes were identified based on the enrichment of DNA barcodes associated with each of the tested peptide specificities (LogFc>2 and p <0.001, using barracoda). Significant T cell recognition of individual peptide sequences is colored based on their protein of origin and segregated based on the origin of samples and diagnosis. All peptides with no significant enrichments are shown as gray dots. C) Bar plot showing the prevalence of epitopes recognized by CD8⁺ T cells across the entire cohort. Bars are colored according to the protein of origin, and novel HPV-derived peptides recognized are indicated with an asterisk (*).

A total of 119 CD8 T cell responses against HPV-derived peptides were detected across the full cohort. These HPV-specific CD8 T cells recognized a total of 109 unique pMHC complexes, which included 94 peptide sequences (Figure 5B). Importantly, we identified 37 HPV-derived peptides (corresponding to 44 unique pMHC complexes) as potential novel HPV epitopes that have not previously been reported to be recognized by T cells (Figure 5C and Supplementary Table 2). In our cohort, HPV-specific CD8 T cells more frequently recognized HPV-derived peptides presented on HLA-A11:01 and HLA-C05:01 (Supplementary Figure 5B). The number of E2 epitopes recognized by these CD8 T cells was higher compared to E6 and E7 epitopes, which was expected given the larger size of E2 proteins. When normalizing for the number of tested peptides, there were no significant differences in the recognition of different HPV proteins by HPV-specific CD8 T cells (Supplementary Figure 5C). T cell recognition to a particular HPV-derived peptide was observed in only few cases across the cohort, reflecting a general low prevalence of CD8 T cell recognition for given pMHC. Only four pMHCs were recognized by CD8 T cells in more than 20% of our cohort (Figure 5C), indicating a high degree of individual variability in the immune response.

### Impaired HPV-specific CD8 T cell response in patients with cervical cancer

To comprehensively understand the CD8 T cell response to HPV, it is essential to evaluate both the breadth and magnitude of their reactivity. The breadth refers to the diversity, or the number of different HPV-specific epitopes recognized by CD8 T cells, while the magnitude describes the frequency of these HPV-specific T cells in the CD8 T cell pool. Our screening results showed a significant difference in the breadth of the HPV-specific CD8 T cell response among the different groups. Specifically, the cancer group exhibited a significantly reduced breadth and lower frequency of these T cells in their blood compared to patients in the CIN3 group and controls (Figure 6A-B). This reduction indicates a lower breadth and frequency of HPV-specific CD8 T cell responses in patients with cancer.

**Figure 6:**
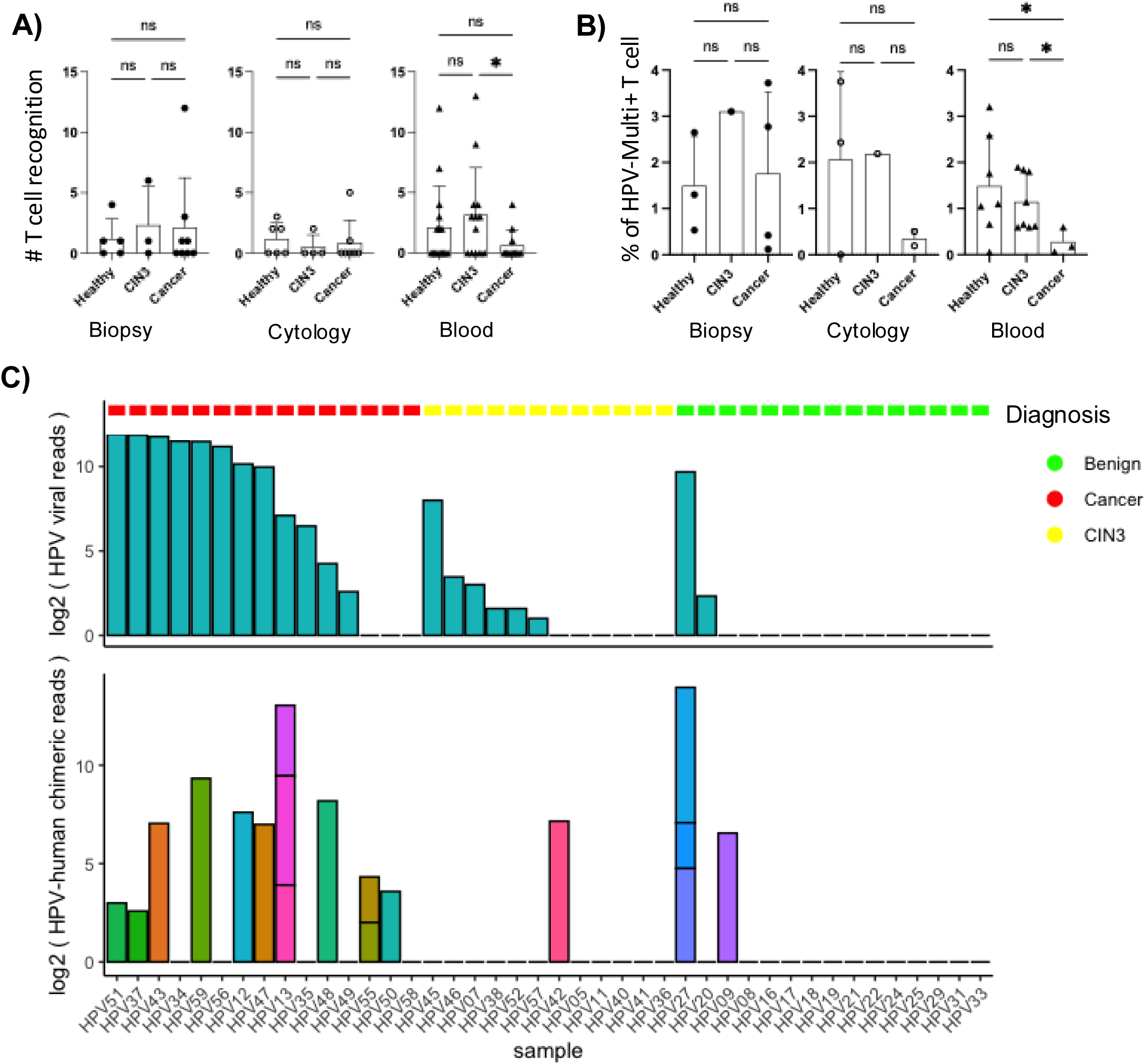
Impaired HPV recognition by CD8 T cells in cancer patients. A) Box plot comparing the number of HPV epitopes recognized by CD8 T cells per patient between control, CIN3, and cancer groups in the biopsy, LBC, and blood. B) Box plot comparing the estimated frequency of recognized HPV epitopes recognized out of total HPV-specific CD8 T cells; between control, CIN3, and cancer groups in the biopsy, LBC, and blood. C) Barplot with the number of reads aligning to HPV genomes per sample, calculated by GATL pathseq. The bottom part shows the number of chimeric reads, that are partially aligned to human and to HPV genomes, calculated by arriba. Plots show mean ± SD (Kruskal-Wallis uncorrected Dunn’s test, *: p<0.05, **: p<0.01, ***: p< 0.001). MFI: mean fluorescence intensity. Filled circles represent values from cervical biopsy samples, open circles from liquid-based cytology (LBC), and triangles from blood.

We observed the presence of HPV-specific CD8 T cell reactivity in healthy control donors and clinically HPV negative CIN3 patients, indicating prior exposure to HPV in these groups. To further investigate the history of HPV infection within our cohort, we analysed RNA sequencing data from cervical samples across all groups. Our RNA sequencing analysis revealed the presence of both HPV viral RNA and evidence of HPV integration into the human genome in all study groups including in some healthy control donors and CIN3 patients (Figure 6C). As expected, the abundance of HPV viral reads and HPV integration was higher in the cancer group compared to the CIN3 and control groups. This finding suggests that HPV infection and integration are not exclusive to individuals with active cervical disease but can also occur in apparently controls.

### Distinct gene expression profiles in cervical cancer compared to CIN3 and normal tissues

Consistent with our earlier findings in this study, Principal Component Analysis (PCA) of RNA sequencing data from cervical biopsies revealed distinct differences between cancer samples and those from CIN3 and controls (Figure 7A). This result illustrated that the gene expression profiles of cancer samples were markedly different from those observed in CIN3 and normal cervical tissues. Interestingly, the cervical samples from healthy controls and CIN3 patients exhibited similar gene expression patterns (Figure 7A). This similarity indicates that normal and CIN3 samples share more similar gene expression profiles, which are distinctly different from those observed in cancer. The heatmap illustrates the differentially expressed genes (DEGs) identified in each group, providing a visual representation of the gene expression variations among controls, CIN3 patients, and cancer patients (Figure 7B). A total of 113 and 13 genes showed higher expression in the normal and CIN3 groups, respectively, compared to the other two groups, while 1263 genes displayed increased expression levels in the cancer group relative to the cervical samples from the CIN3 and control groups (Figure 7B and Supplementary Table 2). To further interpret our RNA-sequencing data, we used the tool developed by Bagaev et al.^18^ to analyse the tumor microenvironment and classify samples based on gene expression profiles (Figure 7C). It was not surprising that the proliferation gene signature score is higher in cervical cancer tissues compared to CIN3 and normal tissues (Figure 7C and Supplementary Figure 6).

**Figure 7:**
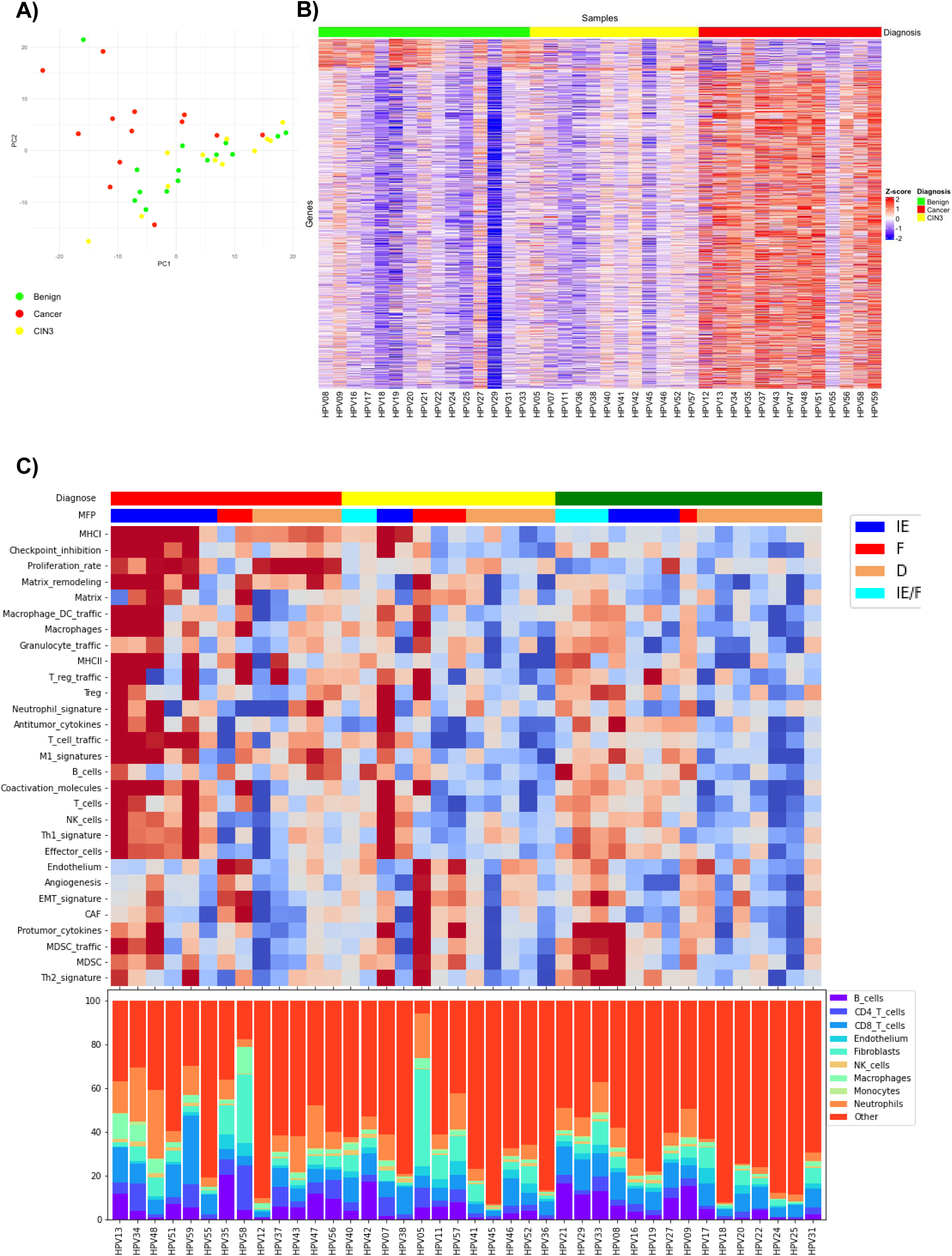
Transcriptome profiling. A) Principal Component Analysis (PCA) performed on the cervical biopsy RNA-seq expression data. Each dot represents a patient and is colored according to the diagnosis: healthy control (green), CIN3 (yellow), and cancer (red). B) Heatmap showing the differentially expressed genes, calculated by DESeq2 with FDR ≤ 0.05 and abs(log2FC) > 1.0 between groups. C) Heatmap of cervical samples from control, CIN3, and cancer groups showing tumor microenvironmental (TME) cell types and biological processed gene signatures, samples ordered by their TME type according to the predictor from Bagaev et al 2022.

Additionally, the gene signature scores for tissue remodelling, antigen presentation via both MHC-I and MHC-II, as well as immune inhibition via checkpoints, were elevated in cervical cancer tissues compared to CIN3 and normal tissues (Figure 7C and Supplementary Figure 6). The microenvironment cell populations-counter (MCP-counter) deconvolution algorithm can analyse the frequency of different cell types in the tissue. Interestingly, our data revealed that the frequency of regulatory T cells is higher in cervical cancer tissue compared to tissues from CIN3 and controls (Figure 7C and Supplementary Figure 7). This finding aligns with our flow cytometry data, which also showed an elevated frequency of Tregs in tumors. Although our flow cytometry data showed a lower percentage of macrophages in the tumors compared to cervical tissues from controls, our MCP-counter deconvolution analysis revealed that the frequency of macrophages, particularly regulatory macrophages (M2 macrophages), is increased in cervical tumor tissues compared to cervical tissue samples from CIN3 and controls (Figure 7C and Supplementary Figure 7).

In summary, our transcriptomic analysis on cervical samples revealed that tumor tissues have distinct gene expression profiles compared to CIN3 and normal tissues, with significant molecular alterations associated with cancer development, including cell proliferation and an immunosuppressive microenvironment. Additionally, the increased frequency of regulatory T cells and regulatory macrophages in cervical cancer tissues highlights the unique immune microenvironment of these tumors.

## Discussion

Our comprehensive analysis of the immune landscape in cervical cancer, CIN3, and normal cervical tissues has revealed critical insights into the immunological changes associated with disease progression. Using high-dimensional flow cytometry analysis, we identified a significant enrichment of late-stage activated or exhausted tumour infiltrating CD8 and CD4 T cells in cervical cancer. Particularly, this exhausted phenotype was largely observed in the cancer group, with no significant differences observed between CIN3 patients and controls. This finding suggests that the emergence of exhausted tumour-infiltrating T cells is associated with the transition from precancerous lesions to cervical cancer. The exhausted CD8 T cell subset (CD39^+^CD103^+^PD-1^+^CD27^+^ CD8 TEM) was enriched both in the tumour biopsies and the LBC samples of cervical cancer patients. This observation aligns with previous studies showing that CD39^+^CD103^+^ CD8 T cells display an exhausted and tissue-resident memory gene signature in various solid tumours including melanoma, lung cancer, head and neck cancer, ovarian cancer, and rectal cancer.^16^ It has been suggested that the presence of CD39^+^CD103^+^ CD8 T cells may serve as a biomarker of potential responsiveness to checkpoint blockade.^19^ The presence of these cells in both tumour biopsies and LBC samples underscores the potential of using LBC samples as a minimally invasive method for studying cervical cancer. The CD57^+^CD39^+^CD103^+^PD-1^+^CD27^+^ CD8 TEM cluster correlated with cancer stage, suggesting that the accumulation of this specific subset may be associated with disease progression. This finding is consistent with previous studies demonstrating the prognostic significance of CD57^+^ CD8 T cells in non-small cell lung cancer.^20,21^

In contrast to CD8 T cells, the triple positive (CD39^+^CD103^+^PD-1^+^) CD4 TEM subset was not enriched in the tumour biopsies compared to non-tumour biopsies. However, the percentage of CD39^+^PD-1^+^CD27^+^ CD4 TEM cells was significantly higher in cancer patients, indicating the presence of an exhausted CD4 T cell population. This exhausted phenotype was further supported by the increased expression of CD39 and EOMES in CD4 T cells from tumour biopsies compared to non-tumour tissues. Interestingly, the frequency of naïve circulating CD8 and CD4 T cells was lower in the cancer group compared to CIN3 and control groups, suggesting a shift towards a more differentiated T cell compartment in cervical cancer patients. This finding is consistent with previous studies demonstrating the depletion of naïve T cells in various cancers.^22^ Altogether, our study provides a comprehensive characterization of the tumour immune landscape in cervical cancer progression, revealing the accumulation of exhausted tumour-infiltrating CD8 and CD4 T cells as a hallmark of the transition from precancerous lesions to invasive cervical cancer. Moreover, our results highlight the importance of analysing the tumour microenvironment directly to capture the most relevant immunological changes, as the phenotype of circulating T cells may not accurately reflect the intratumoral immune status.

Our results also revealed the reduced frequency of cDCs in the peripheral blood and tumour tissues of cervical cancer patients compared to CIN3 and controls. This suggests a reduced presence of cells involved in antigen presentation and initiation of anti-tumour immune responses. This finding is consistent with previous studies demonstrating the importance of cDCs, particularly the cross-presenting cDC1 subset, in orchestrating cytotoxic T cell responses against cancer.^22,23^ In contrast, we observed an increased percentage of pDCs in the cervical tissues of both cancer patients and those with CIN3 lesions compared to controls. pDCs are known for their ability to produce high levels of type I interferons in response to viral infections, which can contribute to anti-HPV immunity. However, in the context of cancer, pDCs have also been shown to promote immune tolerance through the induction of regulatory T cells and the suppression of effector T cell responses.^24,25^ The elevated presence of pDCs in the cervical tissues of cancer and CIN3 patients may therefore reflect a “double-edged sword”, where pDCs contribute to both anti-viral and pro-tumorigenic immune responses.

Interestingly, our analysis of macrophage subsets revealed a significantly lower frequency of CD14+CD64+ macrophages in the cervical tissues of cancer patients compared to controls. This macrophage subset has been associated with a pro-inflammatory phenotype and the ability to present antigens and activate T cells.^26–28^ The reduction in this macrophage population in the tumour microenvironment may further compromise antigen presentation and anti-tumour immunity. In addition, we observed a higher percentage of intermediate monocytes in the peripheral blood of cancer patients compared to healthy controls. Intermediate monocytes have been linked to an immunosuppressive, tumour-promoting phenotype in various cancers.^29,30^ The increased presence of this myeloid subset in the circulation of cervical cancer patients may reflect a systemic shift towards a more immunosuppressive myeloid cell landscape. Taken together, our findings on myeloid cells suggest that the development of cervical cancer is associated with a profound remodelling of the dendritic cell and macrophage compartments, both locally within the tumour microenvironment and systemically in the peripheral blood. The reduction in cDCs and CD14+CD64+ macrophages, associated with the expansion of pDCs and immunosuppressive monocyte subsets, may contribute to the impaired anti-tumour immune responses observed in cervical cancer patients.

The higher expression of PD-L1 on various myeloid cell subsets in the tumour microenvironment further supports the remodelling of the myeloid compartment in cervical cancer. PD-L1 expression is a well-established mechanism of immune evasion, inhibiting T cell function upon binding to PD-1.^31,32^ The elevated PD-L1 expression on macrophages, cDCs, pDCs, and CD11b^+^HLA^-^DR^lo^ cells suggests a broad suppression of anti-tumour immunity within the tumour microenvironment. These findings complement our earlier observations on the altered frequencies of cDCs, pDCs, and macrophage subsets. The increased PD-L1 expression on these cells provides a potential mechanism for their contribution to the immunosuppressive milieu. For instance, the elevated presence of PD-L1 on pDCs suggests skewing towards an immunosuppressive.^24,25^ We also examined CD103 expression in myeloid cell subsets to explore their potential immunosuppressive roles. Our analysis revealed that CD103 expression was elevated in three distinct myeloid populations (including cDCs, CD14^+^CD64^+^ macrophages, and CD11b^-^ HLA^-^DR^Lo^ cells) within cervical tumors compared to tissues from CIN3 patients and controls. While CD103 expression on cDCs is known to promote immunosuppression, the specific role of CD103 expression on macrophages remains unclear. CD103⁺ cDCs have been shown to promote regulatory T cell differentiation and enhance regulatory immune responses in mucosal tissues through retinoic acid and TGF-β.^33,34^

Our findings of increased regulatory T cells and elevated PD-L1 expression on myeloid cells in the tumour microenvironment of cervical cancer patients align with the current understanding of immune evasion mechanisms in cancer disease. The significant rise in Tregs within tumor tissues, as opposed to peripheral blood, highlights the localized nature of immune modulation in cervical cancer. This is in line with studies demonstrating the role of Tregs in suppressing effector T cell responses and promoting immune tolerance in various cancers.^35^ In patients with cervical cancer, the high levels of Treg cells were found to be correlated with shorter survival.^36^ The significant increase in regulatory T cells within tumor tissues, but not in peripheral blood, underscores the localized nature of immune modulation in cervical cancer.

LBC offers a minimally invasive sample source that may permit limited immune-profiling analyses, although its capacity to substitute for cervical biopsies remains to be fully established. Our findings demonstrate that LBC samples reliably capture T cell exhaustion signatures observed in tumor biopsies, supporting their use as a minimally invasive sampling method for monitoring T cell immune responses. This approach offers clear advantages in terms of patient comfort and feasibility for longitudinal studies. However, myeloid populations were markedly underrepresented in LBC compared to biopsies, limiting its utility for comprehensive characterization of innate immune compartments. Despite this limitation, the consistent reduction of conventional dendritic cells (cDCs) in cancer samples across both LBC and biopsy specimens indicates that LBC can partially reflect the cervical immune landscape.

Our study represents an advancement in understanding the CD8 T cell immune response to HPV by extending the focus beyond the well-studied E6 and E7 oncoproteins to include the E2 protein. Screening for T cell responses against E2 protein provides a more comprehensive view of the immune response to HPV, beyond previous focus on only E6 and E7 proteins. T cell responses directed against the E2 protein have been suggested to correlate with HPV clearance and disease regression. One study demonstrated that stimulation with E2 peptides induced T cell proliferation (indicative of a functional memory response) in 50% of healthy women presumed to have cleared HPV16 and in 40% of regressive CIN3 cases.^37^ Here we evaluated the T cell recognition towards all MHC binding ligands from HPV16 and HPV18 proteins E2, E6 and E7, for 14 different HLA alleles. We predicted and screened more than 1100 HLA-binding peptides, ultimately revealing 119 HPV-derived sequences that are recognized by CD8 T cells across our cohort. The inclusion of the E2 protein in our screening is particularly noteworthy. E2, being larger than E6 and E7, accounted for approximately 60% of the total predicted peptides. This highlights its substantial contribution to the HPV antigenic repertoire. Previous studies have largely concentrated on E6 and E7 due to their direct involvement in oncogenesis, but our findings underscore the potential of E2-specific responses in contributing to anti-HPV immunity. Importantly, we reported a total of 37 unique novel epitopes resulting in 44 unique pMHC complexes. This expands our knowledge of sequences in HPV, that are subjected to T cell recognition, in line with findings of La Gruta et al.^38^, who highlighted the importance of broad epitope coverage in understanding the full spectrum of T cell responses.

The presence of tumor infiltrating HPV-specific CD8 T cells has been reported to correlate with improved survival in cervical cancer patients.^39^ Our study highlights a significant impairment in the HPV-specific CD8 T cell response in patients with cervical cancer, characterized by a reduced breadth and frequency of these T cells. This finding aligns with previous research suggesting that the immune system’s ability to recognize and respond to HPV antigens is compromised in head and neck cancer patients.^40^ The observed reduction in immune surveillance may indicate that the immune system’s capacity to control HPV infection and limit cancer progression is significantly compromised in cancer patients.

The HPV infection is widespread and can occur in individuals without clinical symptoms. The infection in many individuals is controlled and ultimately cleared. The presence of HPV-specific CD8 T cell reactivity in healthy control donors is an intriguing observation, suggesting the previous exposure to HPV in these individuals. This is supported by our RNA sequencing data, which revealed HPV viral RNA and evidence of HPV integration in the genomes of healthy controls.

Previous studies have demonstrated distinct molecular signatures between pre-cancerous lesions and cancer tumors.^41,42^ . Our transcriptomic analysis of cervical samples revealed that normal and CIN3 tissues exhibit highly similar gene expression profiles, whereas significant transcriptional changes were observed in cervical cancer. The elevated frequency of regulatory T cells and regulatory macrophages in cervical cancer tissues, as identified by both MCP-counter deconvolution analysis, highlights the suppressive immune landscape of these tumors. These findings are consistent with previous studies that have demonstrated the role of immune cells, especially the regulatory macrophages in promoting tumor growth and evading immune surveillance.^43,44^

In summary, our study provides a comprehensive characterization of the immune landscape in cervical cancer progression. We identified a significant enrichment of exhausted CD8 and CD4 T cells in cervical cancer patients, marking these cells as potential biomarkers for disease progression. Additionally, we observed a reduction in cDCs and macrophages, alongside an increase in pDCs and immunosuppressive monocytes in cancer tissues, indicating a compromised anti-tumor immune response. Elevated PD-L1 expression on myeloid cells and the increased presence of Tregs further support the presence of an immunosuppressive microenvironment within the tumor. Importantly, our findings reveal a reduction in HPV recognition by CD8 T cells in cervical cancer patients. These insights underscore the importance of targeting the immunosuppressive tumor microenvironment, while simultaneously enhancing HPV-specific T cell responses in therapeutic strategies for cervical cancer.

## Materials and Methods

### Sample collection

Clinician collected liquid-based cytology (LBC) specimens, colposcopy guided cervical biopsies (min. 2 x 2 mm), from the cervix and 50 mL peripheral blood was obtained - a full set from each participant. Overall, 57 participants were enrolled, i.e., 25 healthy controls (women without cervical neoplasia), 16 patients with severe neoplasia (CIN 3) and 16 patients with cervical cancer. Some samples had to be excluded due to an insufficient number of cells (Supplementary Table 1). The healthy control donors were recruited from Aleris-Hamlet private hospital, Søborg, Denmark, undergoing HPV unrelated hysterectomy. Specimens were obtained during already planned surgery to minimize discomfort for the patient. Patients with CIN3 were recruited from Department of Gynecology and Obstetrics, Hvidovre Hospital, Hvidovre. Biopsies and cytology specimens were collected prior to having a cone biopsy, to limit blood contamination into the specimen. If possible, the blood samples were taken during that same consultation. Patients with cervical cancer were recruited from Department of Gynecology and Obstetrics, Odense University Hospital, Odense. Specimens were collected prior to assessment of the cancer stage in full anaesthesia. Patients were enrolled based on the following criteria: Inclusion criteria: Female participants, >18 years at inclusion and with informed consent. Exclusion criteria: Patients receiving immunosuppressive treatment, e.g., larger doses of prednisolone (>5mg/ day), previous cancer of any kind, healthy controls with previous cervical neoplasia. Clinical parameters e.g., diagnosis, other diseases (especially immune mediated diseases), medication, age, health status, history of smoking, and other environmental factors was also registered.

All specimens were obtained anonymously, marked with a study number, delivered to the laboratory at the Technical University of Denmark (DTU), Lyngby, Denmark and were all handled within 6 hours after sampling. In total, 11 different types of HPV (16, 18, 33, 39, 45, 51, 52, 53, 61, 70, 82) were detected among all participants (Table S3), 17% being positive for >1 type. Out of these subtypes, HPV 53, 61 and 82 are not classified as high-risk HPV types associated with cervical cancer. 70% of the healthy controls was tested negative for HPV, whereas the vast majority of CIN 3 (90%) and cancer patients (93%) were HPV positive (Table S3). The cancer patients had the highest prevalence of HPV infection, 60% being positive for HPV 16 and HPV 18 was only detected in few of the CIN3 and cancer patients. The cervical cancer patients were classified by disease progression according to the FIGO grading scale^45^ and 87% had invasive stage II or higher (Table S3). 73.3% was squamous cell carcinomas, 20% was adenocarcinomas, one adeno-squamous and adenocarcinoma of mucinous type. BMI was non-significant; (healthy controls: 27 ± 4), (CIN 3: 25 ± 5) and (cancer: 29 ± 10).

### Cell isolation

Liquid-based cytology (LBC) specimens: The LBC specimens were collected using two Cervix-Brush (Rovers) technique. The first sample was kept on ice in 15 mL digestion buffer (1:10 Hank’s Balanced Saline Solution (HBSS) 50X + 15mM 1,5% 4-(2-hydroxyethyl)-1-piperazineethanesulfonic acid (HEPES)) until cell isolation. To isolate the cells, the brush was rinsed, and suspension filtered through a cell strainer. Cells were centrifuged and resuspended in phosphate saline buffer (PBS), counted on NucleoCounter SCC-100TM (Chemometec), using SCC-CassetteTM. Cells were again centrifuged and resuspended in 1 mL of freezing media 10% DMSO (Dimethyl Sulfoxide) Hybrid-Max (Sigma-Aldrich) and 90% fetal calf serum (FCS) (GibcoTM qualified, New Zealand). 5-10 x10^6^ cells/vial and distributed in cryotubes. The vials were frozen by 1°C/min in freezing boxes placed at -80°C and next day transferred to a -180°C nitrogen tank for long-term storage until used for further analysis. The second specimen were kept at room temperature (RT) in 10 mL SurePathTM Collection Vial (BD). These specimens were then sent to Department of Pathology, Hvidovre Hospital, Denmark, for HPV typing using Anyplex II HPV28 detection real-time PCR.

Biopsy: One biopsy from each patient was preserved in RNALater for RNA sequencing. The other biopsy samples were collected in the same digestion buffer as cytology and stored on ice. Using scalpel and tweezer, the biopsies were cut into smaller pieces, centrifuged for homogenization for 1 min using gentleMAC Dissociator (Miltenyi Biotec). 2,5 mL (2U/ml) DNase bovine pancreas (Sigma-Aldrich, Merck) was added to the specimen. After 1-hour incubation (37°C), the specimens were further dissociated (1 min) using gentleMACS. The suspension was then filtered through a cell strainer, centrifuged, resuspended, and counted. From this step the cells were treated as described for the cytology.

PBMC: Peripheral blood samples were collected in five 10 mL Vacutainer heparin tubes and kept at RT until separated using filter tubes (Falcon Leucosep) saturated with 15 mL LymphoprepTM 1.077g/mL, (STEMCELL, cat. 07851), and PBS. After separation, lymphocytes were resuspended and counted on NucleoCounter (Chemometec) and were treated as described for the cytology and biopsy specimens.

### Immunophenotyping using flow cytometry

Cell specimens were stained with the surface antibodies diluted in equal amounts of FACS buffer and Brilliant Stain Buffer (BD) giving a total of 50 μl per sample. Samples were then fixed and permeabilized using fixation/permeabilization kit (Invitrogen). Intracellular antibodies were diluted in washing buffer for intracellular staining for transcription factors and intracellular cytokines. The near-IR viability dye (Invitrogen) was used to discriminate between live and dead cells. Flow cytometry experiments were performed on LSR-Fortessa (BD Biosciences). Data was analysed in FACSDiva software (BD Biosciences) and FlowJo v10.7 (TreeStar, Inc.).

### Statistical analyses

Uniform manifold approximation and projections (UMAP) were made in FlowJo using the UMAP plugin a dimensionality reduction technique. DownSampleV3 was applied on the specimen before FlowSOM was used on concatenated files (clustering and visualization algorithm) to analyse and detect data subsets using self-organizing maps. All plugins for FlowJo from flowjo.com/exchange. Data was analysed with a non-parametric Kruskal-Wallis test with Dunn’s correlation for multiple comparisons. These statistical analyses were conducted using GraphPad Prism 9.0 and later.

### HPV peptides selection

The protein sequence of HPV 16 E2 (ID P03120), E6 (P03126) and E7 (P03129) and HPV 18 E2 (P06790), E6 (P06463) and E7 (P06788) was derived from "http://www.uniprot.org", and MHC-I binding peptides within these proteins were predicted using netMHCpan 4.0^46^ The 14 most common human leucocyte antigen (HLA) were chosen, HLA-A (A01:01, A02:01, A03:01, A11:01, A24:02), HLA-B (B07:02, B08:01, B15:01, B35:01) and HLA-C (C03:04, C04:01, C05:01, C07:01, C07:02). The search criteria in the artificial neural network netMHCpan 4.0 were: 8-11 amino acid in peptide length, Eluted ligand rank (EL rank) >2. In total 1161 potential distinct HLA-binding peptides were selected. Furthermore IEDB.org was used to include additional T cell epitopes previously identified. Search criteria were ≥1 references and ≥2 assays and maximum peptide length of 14. This listed 7 additional peptides. A PubMed search for known CD8 T cell epitopes was also performed and resulted 3 more peptides, added to the screening library. All predicted peptides were custom synthesized with an estimated purity of 70-92% by Pepscan Presto BV, Lelystad, The Netherlands. This generated in total 1238 peptide-HLA pairs for experimental evaluation. The 14 different HLAs were produced using plasmids in Escherichia coli as previously discribed.^47^

### CD8 T cell epitope screening using DNA-barcoded peptide-MHC multimers

The CD8 T cell recognition was evaluated using DNA-barcoded peptide-MHC multimers, covering the full library of predicted HPV-derived MHC binding peptide sequences. ^48^ Briefly, the HPV peptide multimers were generated by assembling unique DNA barcodes and peptide-MHCs on dextran backbones (Fina Biosolutions, Rockville, MD, USA) by a biotin-streptavidin interaction. The biological samples were then stained with DNA-barcoded peptide-MHC multimers and antibodies to determine the CD8 T cell population (show gating strategy in suppl). The pMHC multimer binding CD8 T cells were sorted and the DNA barcodes on the sorted cells were amplified. The amplified DNA were sequenced, and the data analysed using Barracoda (https://services.healthtech.dtu.dk/services/Barracoda-1.8/). The enriched DNA sequenced with LogFc >2 and P < 0.001 indicate the p-MHC recognized by CD8 T cells.

### Transcriptome profiling

The biopsy samples preserved in RNALater were sent to Novogene in UK for RNA sequencing. RNA-seq quality control was perfomed with fastqc wrapper in trim_galore 0.6.4. Gene expression quantification into TPM was performed with Kallisto 0.42.1. RNA-seq bam file was generated by STAR 2.7.9. The principal component analysis (PCA) performed on the cervical biopsy RNA-seq expression data. We calculated the differentially expressed genes for each group, calculated by DESeq2 with FDR ≤ 0.05 and abs (log2FC) > 1.0 between groups. We next did the tumor microenvironment (TME) typing on RNA-seq samples. TMP matrix was log2 normalised, median centered and mad scaled to obtain z-score matrix. Z-score matrix was subjected to TME prediction via KNeighborsClusterClassifier, with pancancer TCGA reference and z-score matrix provided by Bagaev et al. 2021^18^ on the github page: https://github.com/BostonGene/MFP.

Viral integration sites were called with arriba 2.3.0, run on a combination of human and HPV16 and HPV18 viral references. Viral genomes were manually downloaded as fasta and gtf files from NCBI database and united with human reference and STAR genomeGenerate command was ran to create a combined reference. Quantification of viral reads was performed with Pathseq program from gatk/4.0.1.1 toolkit.

### Ethics statement

This study was conducted in accordance with the principles outlined in the Declaration of Helsinki. Approval for the study design and sample collection was obtained from the Committee on Health Research Ethics in the Capital Region of Denmark. All included participants gave informed written consent for inclusion. All specimens were kept anonymously. The study was approval by The Danish Data Protection Agency.

## Acknowledgments

This research was supported by the Danish Cancer Society Research Center (R146-A9531 and R147-A10025) and the Aleris-Hamlet Research Foundation; European Union’s Horizon 2020 research and innovation programme under grant agreement No ImmunoSABR (733008); and MK received funding from the European Union’s Horizon 2020 research and innovation program under the Marie Sklodowska-Curie grant agreement no. 713683 (COFUNDfellowsDTU). We would also like to extend our gratitude to the patients who participated in this study. Additionally, we wish to thank the medical staff at Hvidovre Hospital and Odense University Hospital for their assistance in patient recruitment and sample collection. We also acknowledge the technical and academic staff at the Technical University of Denmark (DTU) for assistance in this research.

---

**Supplementary figure 1:**
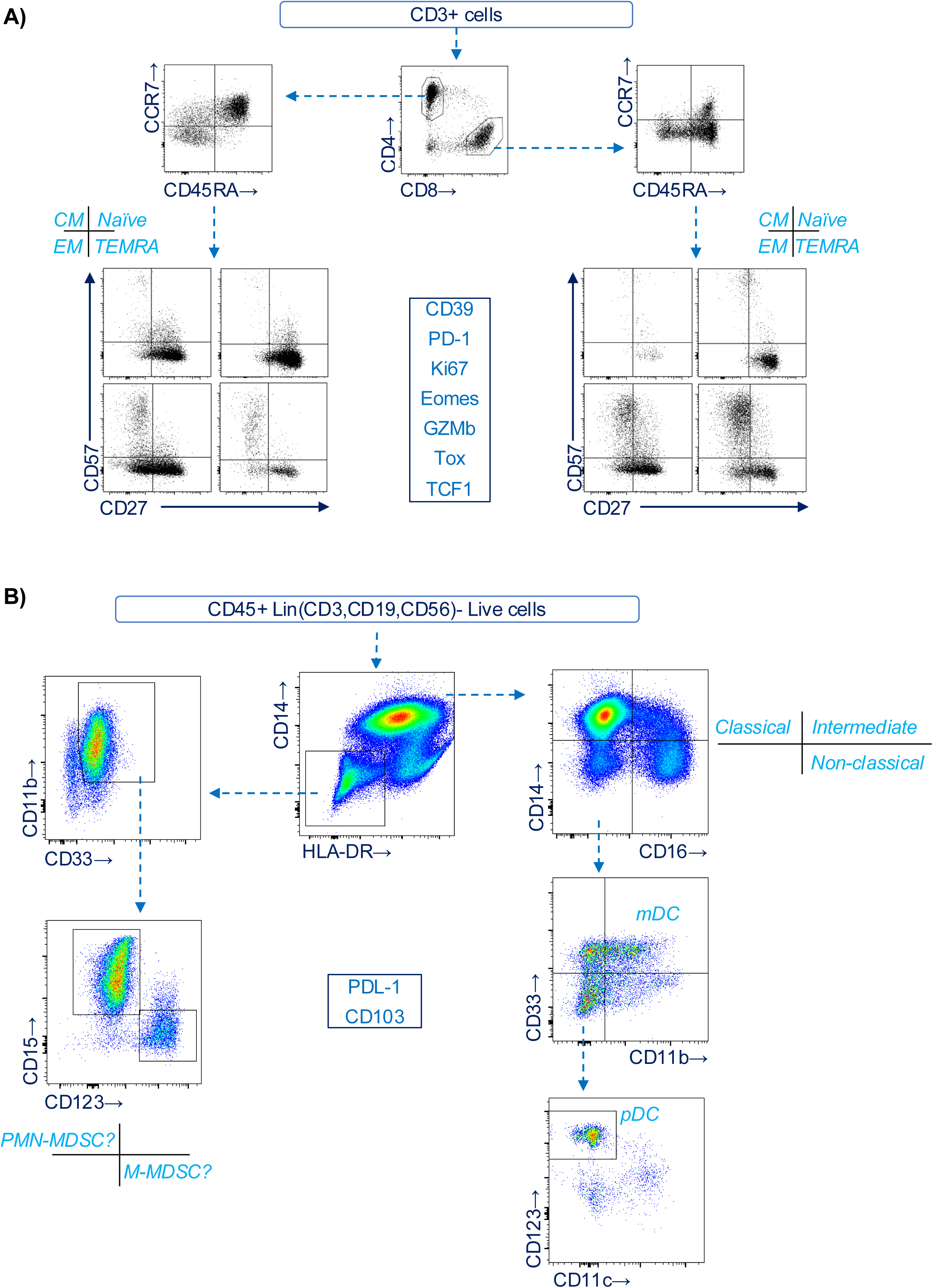
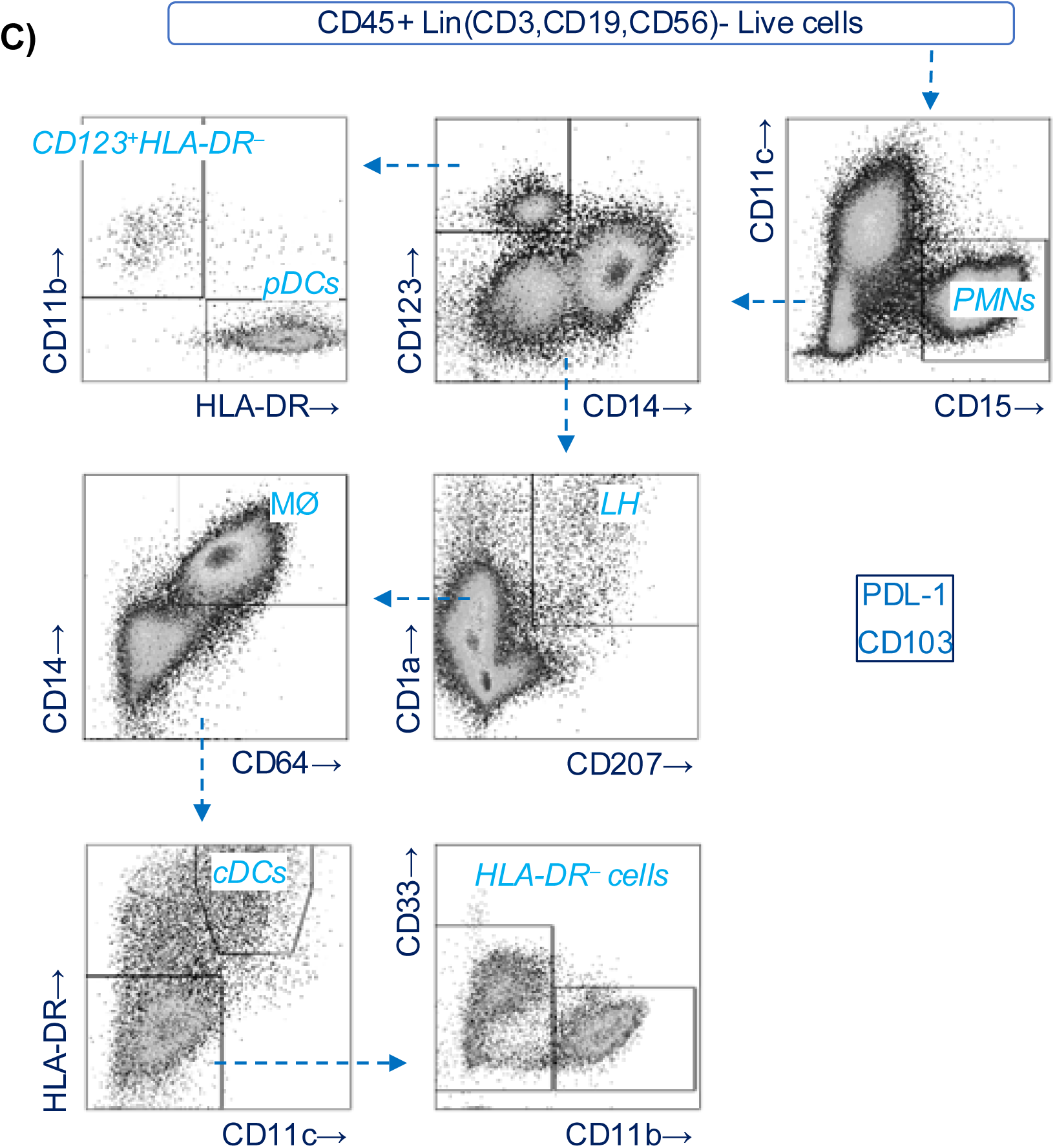
Gating strategies. The gating strategies that were used to identify T cell and myeloid cell subsets in the blood and cervical tissues. T cell subsets were analyzed for the expression of CD39, PD-1, Ki67, Eomes, GZMB, Tox, and TCF-1. Myeloid cell subsets were analyzed for the expression of PDL-1 and CD103.

**Supplementary figure 2:**
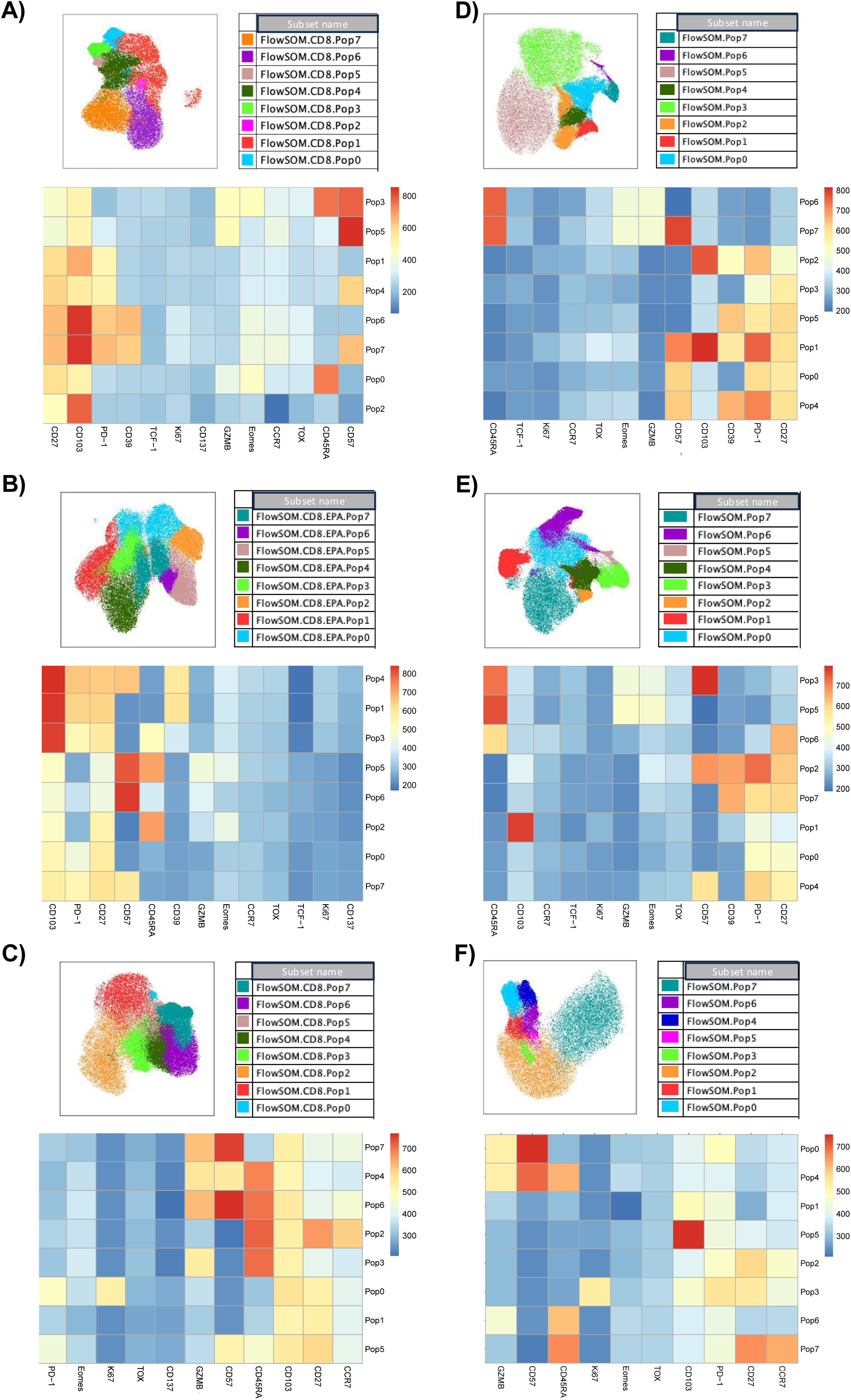
Presence of exhaustion signature in tumor infiltrating CD8 T Cells. UMAP illustration of CD8 T cells in control, CIN3, and cancer groups. CD8 T cells were clustered into 8 different subpopulations using FlowSOM. The heatmap shows the expression level of each marker in different clusters.

**Supplementary figure 3:**
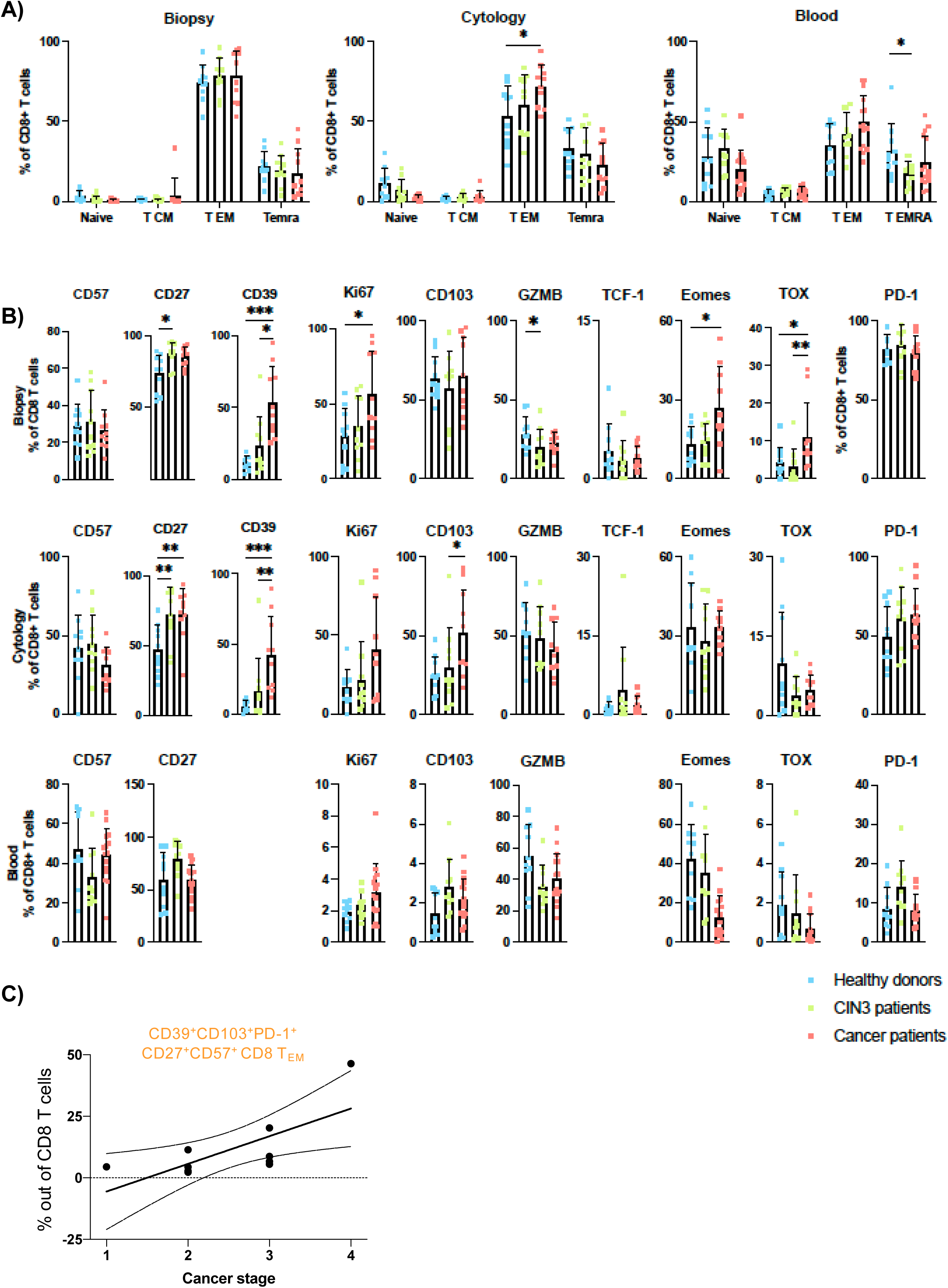
CD8 T cell phenotypes in the biopsy, cytology, and blood samples. A) The percentage of naïve, TCM, TEM, TEMRA CD8 T cell subsets in biopsy, cytology, and blood of healthy control donors, CIN3 patients and cancer patients. B) The percentage of indicated marker expressing CD8 T cells in biopsy, cytology, and blood of healthy control donors, CIN3 patients and cancer patients. Plots show mean ± SD (Kruskal-Wallis uncorrected Dunn’s test, *: p<0.05, **: p<0.01, ***: p< 0.001).

**Supplementary figure 4:**
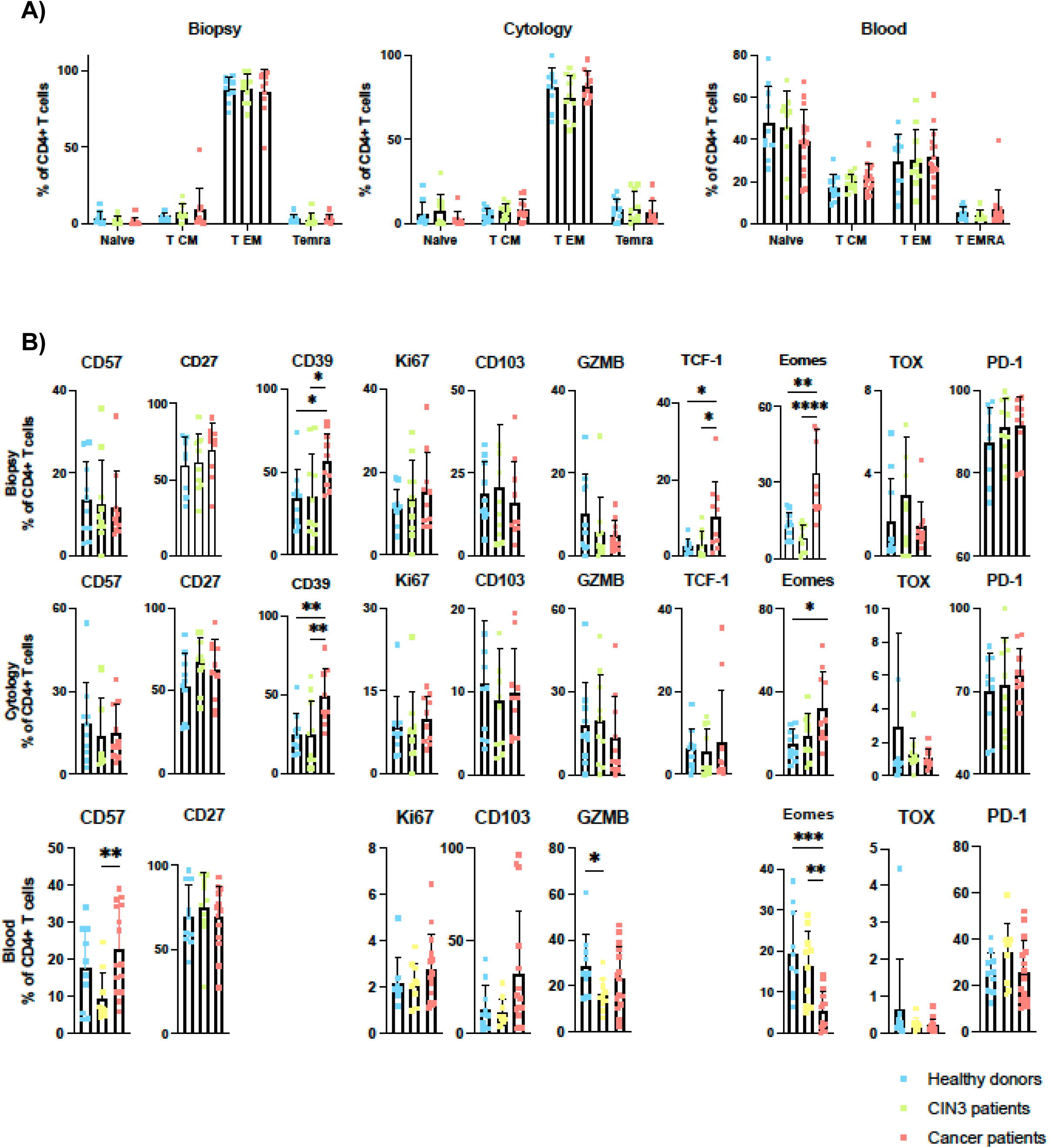
CD4 T cell phenotypes in the biopsy, cytology, and blood samples. A) The percentage of naïve, TCM, TEM, TEMRA CD4 T cell subsets in biopsy, cytology, and blood of healthy control donors, CIN3 patients and cancer patients. B) The percentage of indicated marker expressing CD4 T cells in biopsy, cytology, and blood of healthy control donors, CIN3 patients and cancer patients. Plots show mean ± SD (Kruskal-Wallis uncorrected Dunn’s test, *: p<0.05, **: p<0.01, ***: p< 0.001).

**Supplementary figure 5:**
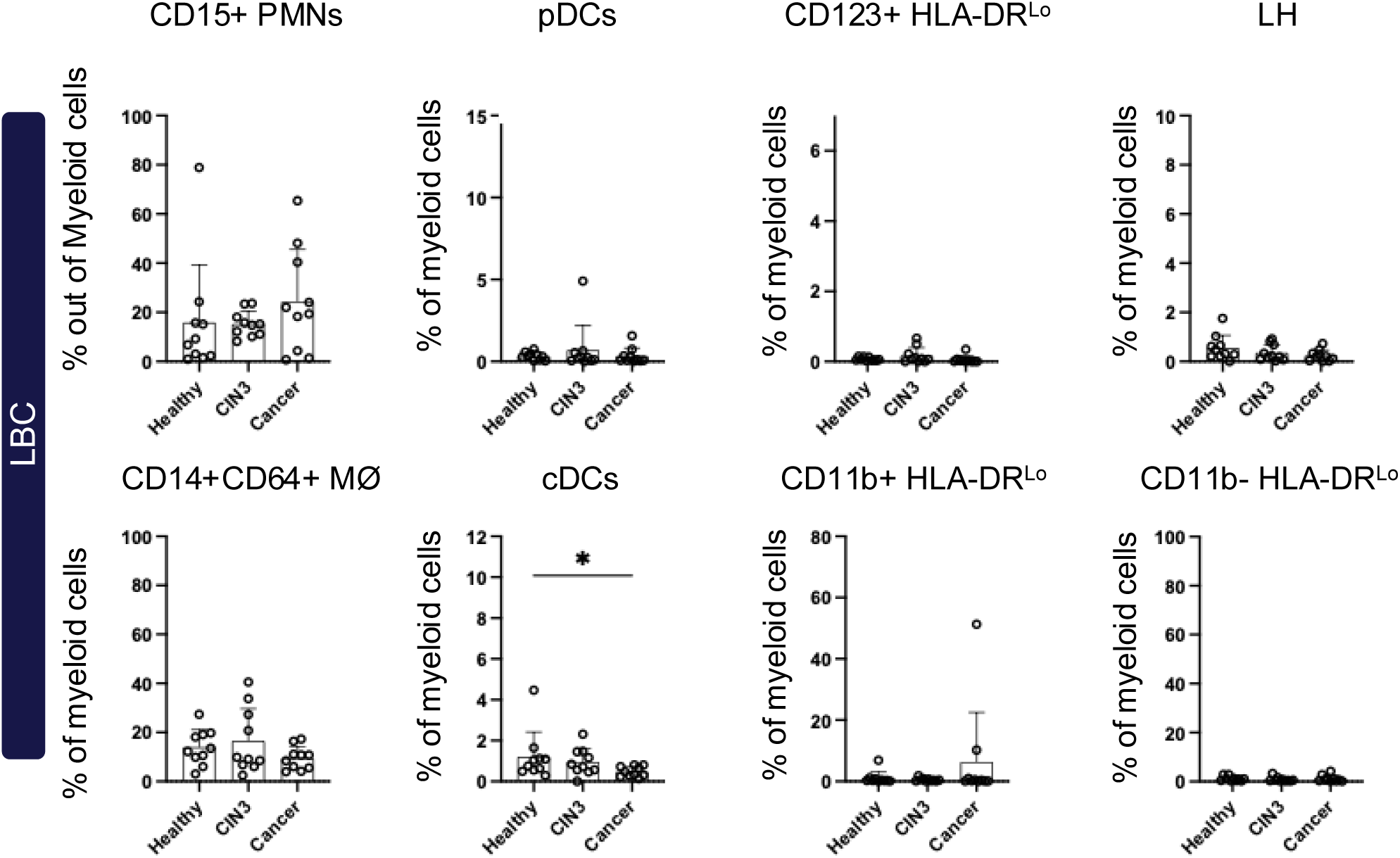
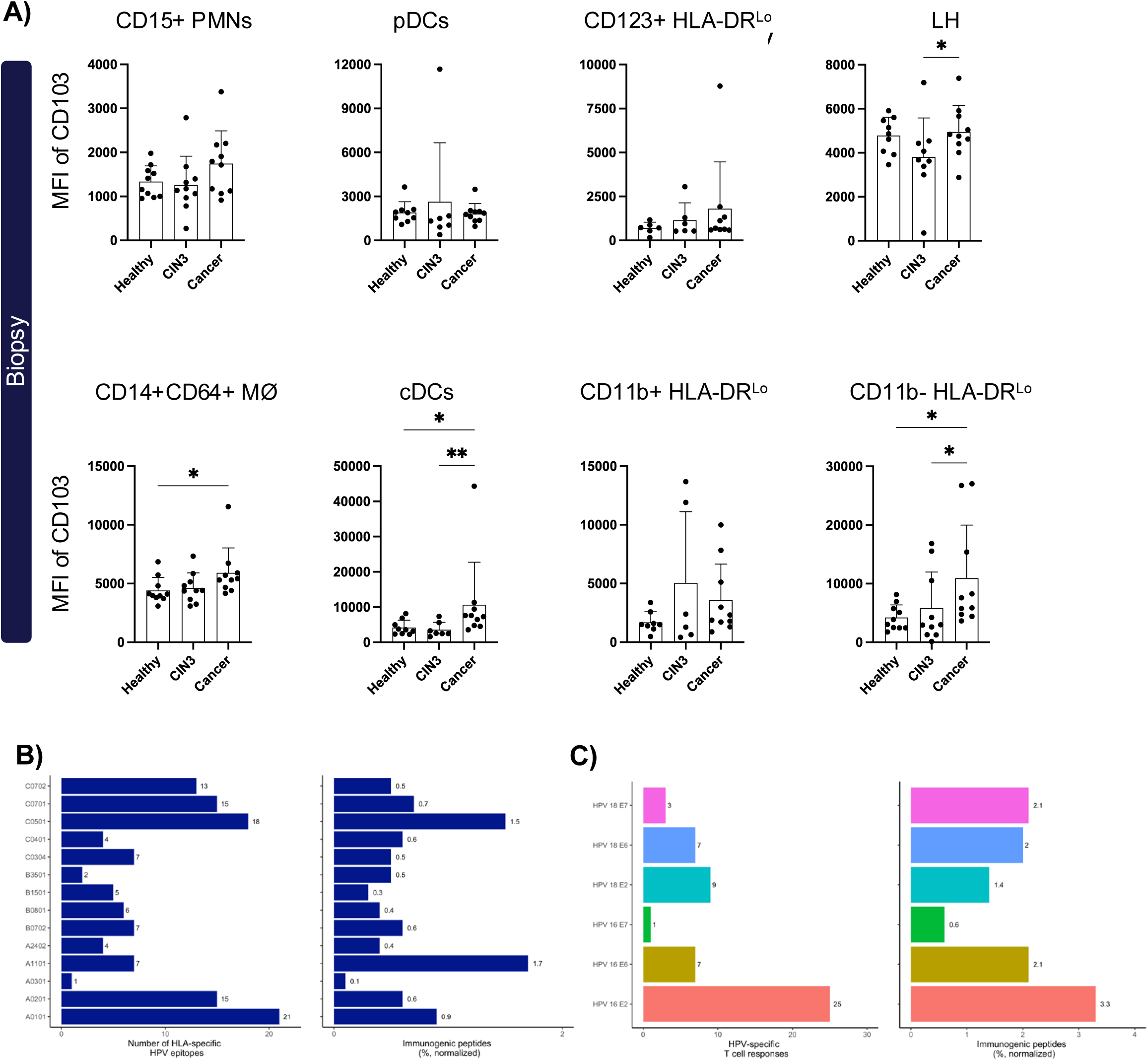
CD103 expression on subset of tissue myeloid cells, and prevalence of recognized pMHC in our cohort. A) The expression (MFI) of CD103 on different subsets of cervical myeloid cells from biopsy samples is shown. Plots show mean ± SD (Kruskal-Wallis uncorrected Dunn’s test, *: p<0.05, **: p<0.01, ***: p< 0.001). B) Bar plots summarize the number of Number of recognized HPV epitopes by CD8 T cells per HLA and the HLA-restricted recognition (% recognized peptides) in the analyzed patients. Normalized here represents the fraction of T cell recognized peptides out of the total number of peptides analyzed for a given HLA restriction across the HLA-matching donors (% normalized). C) Bar plots show the number of epitopes derived from each of the HPV proteins and their recognition score (% recognized peptides).

**Supplementary figure 6:**
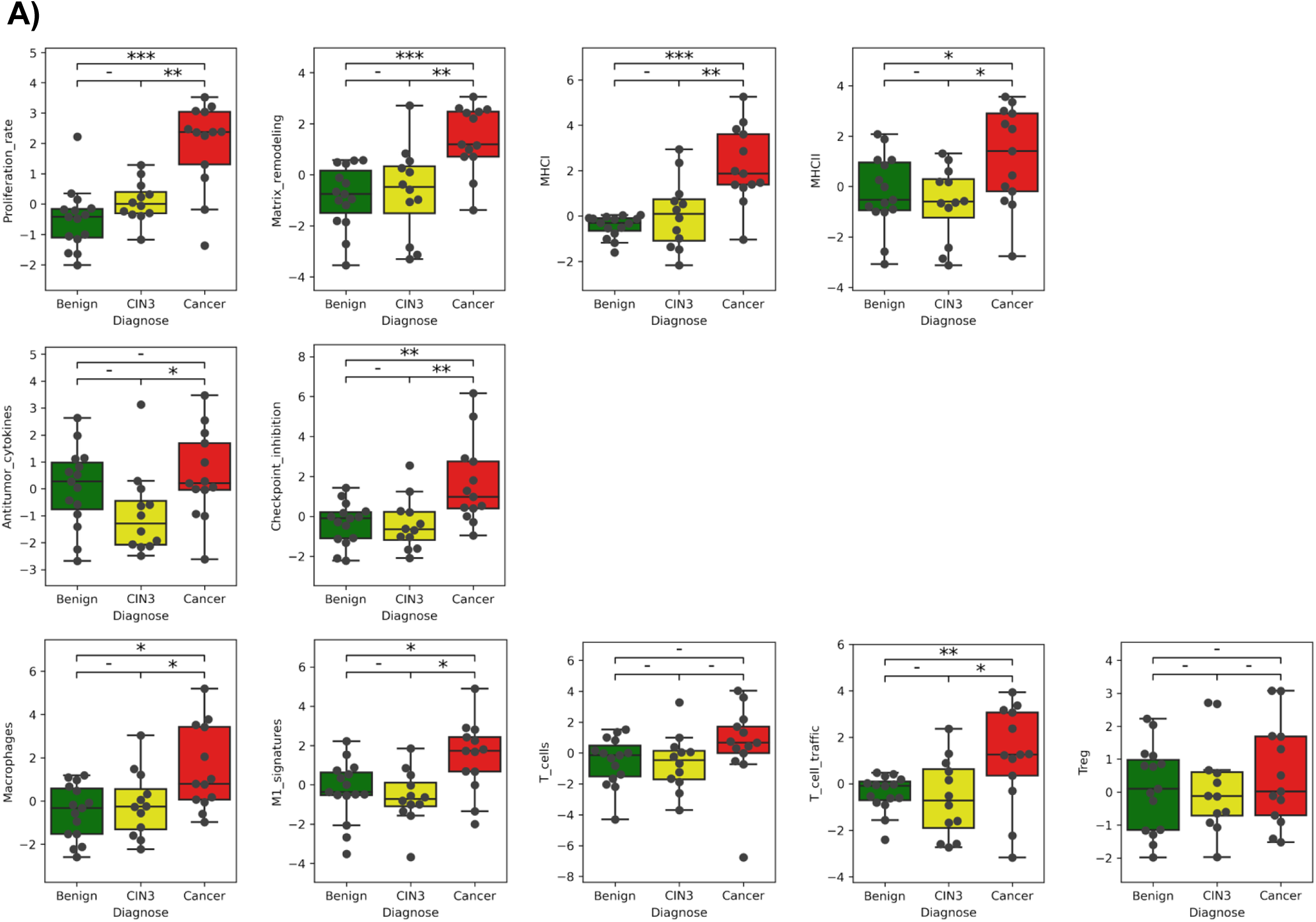
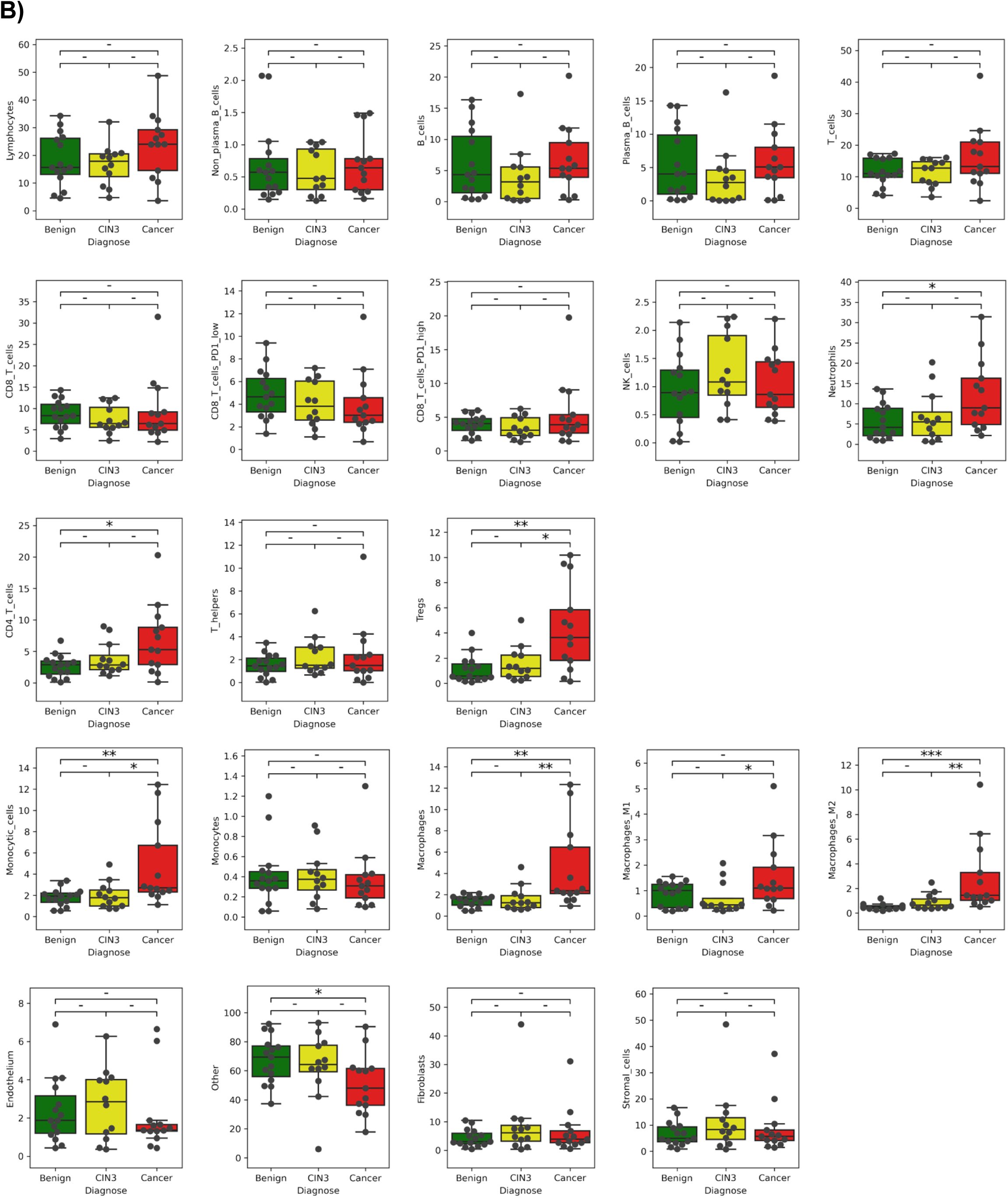
The gene signature scores of cervical samples. Boxplots of gene signature scores of cervical samples from controls, CIN3, and cancer groups. Plots show median ± 25^th^ percentile (Mann-Whitney test, *: p<0.05, **: p<0.01, ***: p< 0.001).

**Supplementary figure 7:** The microenvironment cell populations-counter (MCP-counter) deconvolution analysis. Boxplots of microenvironmental (TME) cell types using microenvironment cell populations-counter (MCP-counter) deconvolution algorithm. Plots show median ± 25^th^ percentile (Mann-Whitney test, *: p<0.05, **: p<0.01, ***: p< 0.001).

**Supplementary Table 1.**
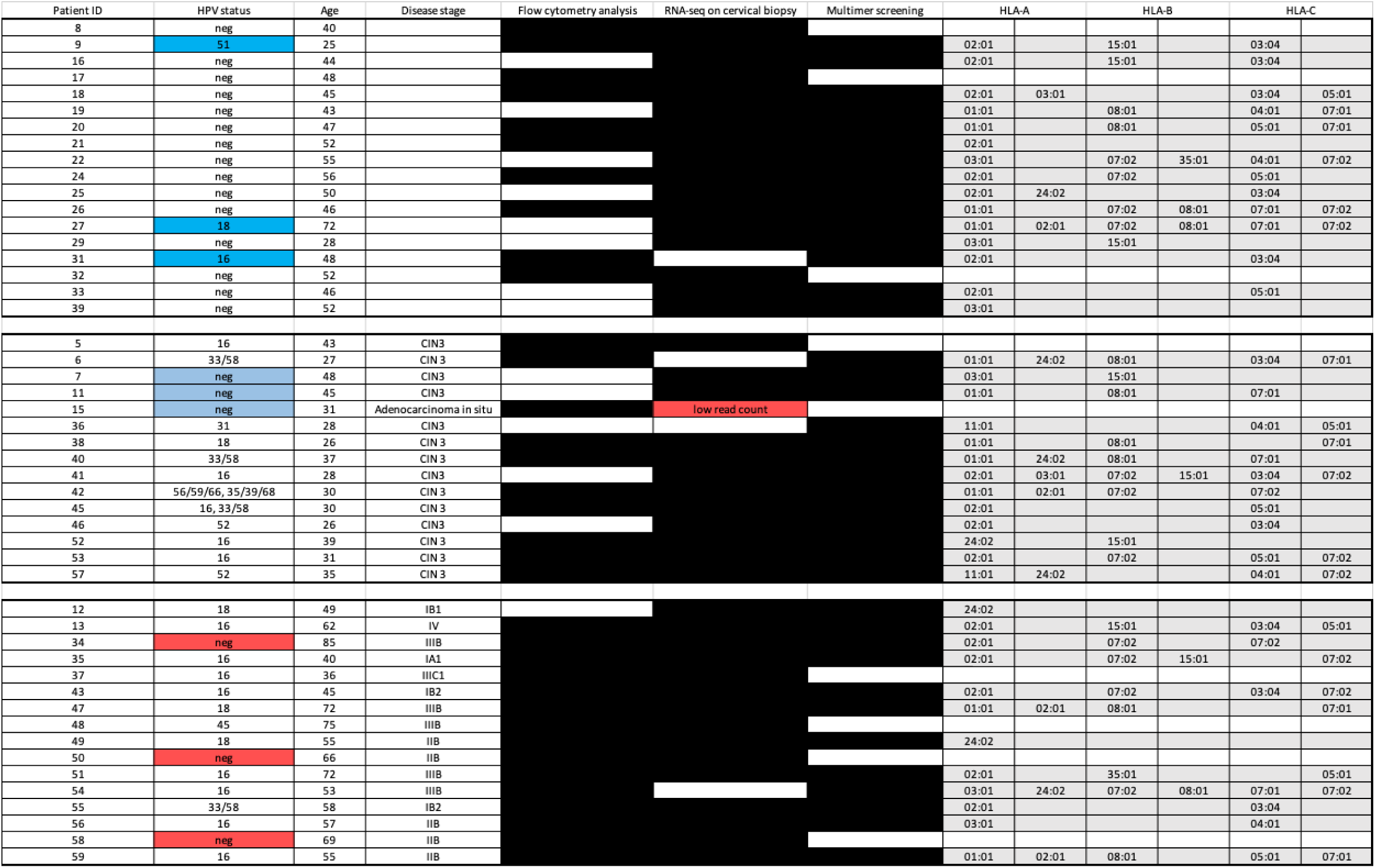

| Patient ID | HPV status | Age | Disease stage | Flow cytometry analysis | RNA-seq on cervical biopsy | Multimer screening | HLA-A |  | HLA-B |  | HLA-C |  |
| --- | --- | --- | --- | --- | --- | --- | --- | --- | --- | --- | --- | --- |
| 8 | neg | 40 |  |  |  |  |  |  |  |  |  |  |
| 9 | 51 | 25 |  |  |  |  | 02:01 |  | 15:01 |  | 03:04 |  |
| 16 | neg | 44 |  |  |  |  | 02:01 |  | 15:01 |  | 03:04 |  |
| 17 | neg | 48 |  |  |  |  |  |  |  |  |  |  |
| 18 | neg | 45 |  |  |  |  | 02:01 | 03:01 |  |  | 03:04 | 05:01 |
| 19 | neg | 43 |  |  |  |  | 01:01 |  | 08:01 |  | 04:01 | 07:01 |
| 20 | neg | 47 |  |  |  |  | 01:01 |  | 08:01 |  | 05:01 | 07:01 |
| 21 | neg | 52 |  |  |  |  | 02:01 |  |  |  |  |  |
| 22 | neg | 55 |  |  |  |  | 03:01 |  | 07:02 | 35:01 | 04:01 | 07:02 |
| 24 | neg | 56 |  |  |  |  | 02:01 |  | 07:02 |  | 05:01 |  |
| 25 | neg | 50 |  |  |  |  | 02:01 | 24:02 |  |  | 03:04 |  |
| 26 | neg | 46 |  |  |  |  | 01:01 |  | 07:02 | 08:01 | 07:01 | 07:02 |
| 27 | 18 | 72 |  |  |  |  | 01:01 | 02:01 | 07:02 | 08:01 | 07:01 | 07:02 |
| 29 | neg | 28 |  |  |  |  | 03:01 |  | 15:01 |  |  |  |
| 31 | 16 | 48 |  |  |  |  | 02:01 |  |  |  | 03:04 |  |
| 32 | neg | 52 |  |  |  |  |  |  |  |  |  |  |
| 33 | neg | 46 |  |  |  |  | 02:01 |  |  |  | 05:01 |  |
| 39 | neg | 52 |  |  |  |  | 03:01 |  |  |  |  |  |
| 5 | 16 | 43 | CIN3 |  |  |  |  |  |  |  |  |  |
| 6 | 33/58 | 27 | CIN 3 |  |  |  | 01:01 | 24:02 | 08:01 |  | 03:04 | 07:01 |
| 7 | neg | 48 | CIN3 |  |  |  | 03:01 |  | 15:01 |  |  |  |
| 11 | neg | 45 | CIN3 |  |  |  | 01:01 |  | 08:01 |  | 07:01 |  |
| 15 | neg | 31 | Adenocarcinoma in situ |  | low read count |  |  |  |  |  |  |  |
| 36 | 31 | 28 | CIN3 |  |  |  | 11:01 |  |  |  | 04:01 | 05:01 |
| 38 | 18 | 26 | CIN 3 |  |  |  | 01:01 |  | 08:01 |  |  | 07:01 |
| 40 | 33/58 | 37 | CIN 3 |  |  |  | 01:01 | 24:02 | 08:01 |  | 07:01 |  |
| 41 | 16 | 28 | CIN3 |  |  |  | 02:01 | 03:01 | 07:02 | 15:01 | 03:04 | 07:02 |
| 42 | 56/59/66, 35/39/68 | 30 | CIN 3 |  |  |  | 01:01 | 02:01 | 07:02 |  | 07:02 |  |
| 45 | 16, 33/58 | 30 | CIN 3 |  |  |  | 02:01 |  |  |  | 05:01 |  |
| 46 | 52 | 26 | CIN3 |  |  |  | 02:01 |  |  |  | 03:04 |  |
| 52 | 16 | 39 | CIN 3 |  |  |  | 24:02 |  | 15:01 |  |  |  |
| 53 | 16 | 31 | CIN 3 |  |  |  | 02:01 |  | 07:02 |  | 05:01 | 07:02 |
| 57 | 52 | 35 | CIN 3 |  |  |  | 11:01 | 24:02 |  |  | 04:01 | 07:02 |
| 12 | 18 | 49 | IB1 |  |  |  | 24:02 |  |  |  |  |  |
| 13 | 16 | 62 | IV |  |  |  | 02:01 |  | 15:01 |  | 03:04 | 05:01 |
| 34 | neg | 85 | IIIB |  |  |  | 02:01 |  | 07:02 |  | 07:02 |  |
| 35 | 16 | 40 | IA1 |  |  |  | 02:01 |  | 07:02 | 15:01 |  | 07:02 |
| 37 | 16 | 36 | IIIC1 |  |  |  |  |  |  |  |  |  |
| 43 | 16 | 45 | IB2 |  |  |  | 02:01 |  | 07:02 |  | 03:04 | 07:02 |
| 47 | 18 | 72 | IIIB |  |  |  | 01:01 | 02:01 | 08:01 |  |  | 07:01 |
| 48 | 45 | 75 | IIIB |  |  |  |  |  |  |  |  |  |
| 49 | 18 | 55 | IIB |  |  |  | 24:02 |  |  |  |  |  |
| 50 | neg | 66 | IIB |  |  |  |  |  |  |  |  |  |
| 51 | 16 | 72 | IIIB |  |  |  | 02:01 |  | 35:01 |  |  | 05:01 |
| 54 | 16 | 53 | IIIB |  |  |  | 03:01 | 24:02 | 07:02 | 08:01 | 07:01 | 07:02 |
| 55 | 33/58 | 58 | IB2 |  |  |  | 02:01 |  |  |  | 03:04 |  |
| 56 | 16 | 57 | IIB |  |  |  | 03:01 |  |  |  | 04:01 |  |
| 58 | neg | 69 | IIB |  |  |  |  |  |  |  |  |  |
| 59 | 16 | 55 | IIB |  |  |  | 01:01 | 02:01 | 08:01 |  | 05:01 | 07:01 |

## References

1. WHO Cervical Cancer. https://www.who.int/news-room/fact-sheets/detail/cervical-cancer https://www.who.int/news-room/fact-sheets/detail/cervical-cancer (2024).

2. Molijn, A. et al. The complex relationship between human papillomavirus and cervical adenocarcinoma. Int J Cancer 138, (2016).

3. International Agency for Research on Cancer. Cervix uteri Source: Globocan 2020. Globocan 419, (2020).

4. Small, W., et al. Cervical cancer: A global health crisis. Cancer vol. 123 Preprint at 10.1002/cncr.30667 (2017).

5. Shanmugasundaram, S. & You, J. Targeting persistent human papillomavirus infection. Viruses vol. 9 Preprint at 10.3390/v9080229 (2017).

6. McCredie, M. R. et al. Natural history of cervical neoplasia and risk of invasive cancer in women with cervical intraepithelial neoplasia 3: a retrospective cohort study. Lancet Oncol 9, (2008).

7. Sabatini, M. E. & Chiocca, S. Human papillomavirus as a driver of head and neck cancers. British Journal of Cancer vol. 122 Preprint at 10.1038/s41416-019-0602-7 (2020).

8. Strander, B., Andersson-Ellström, A., Milsom, I. & Sparén, P. Long term risk of invasive cancer after treatment for cervical intraepithelial neoplasia grade 3: Population based cohort study. Br Med J 335, (2007).

9. Wei, F., Georges, D., Man, I., Baussano, I. & Clifford, G. M. Causal attribution of human papillomavirus genotypes to invasive cervical cancer worldwide: a systematic analysis of the global literature. The Lancet 404, (2024).

10. de Sanjose, S. et al. Human papillomavirus genotype attribution in invasive cervical cancer: a retrospective cross-sectional worldwide study. Lancet Oncol 11, (2010).

11. Melsheimer, P., Vinokurova, S., Wentzensen, N., Bastert, G. & Von Knebel Doeberitz, M. DNA Aneuploidy and Integration of Human Papillomavirus Type 16 E6/E7 Oncogenes in Intraepithelial Neoplasia and Invasive Squamous Cell Carcinoma of the Cervix Uteri. Clinical Cancer Research 10, (2004).

12. von Knebel Doeberitz, M., Rittmüller, C., Aengeneyndt, F., Jansen-Dürr, P. & Spitkovsky, D. Reversible repression of papillomavirus oncogene expression in cervical carcinoma cells: consequences for the phenotype and E6-p53 and E7-pRB interactions. J Virol 68, (1994).

13. Doorbar, J. Model systems of human papillomavirus-associated disease. Journal of Pathology vol. 238 Preprint at 10.1002/path.4656 (2016).

14. Ghanaat, M. et al. Virus against virus: strategies for using adenovirus vectors in the treatment of HPV-induced cervical cancer. Acta Pharmacologica Sinica vol. 42 Preprint at 10.1038/s41401-021-00616-5 (2021).

15. Sharma, P. & Allison, J. P. The future of immune checkpoint therapy. Science (1979) 348, (2015).

16. Duhen, T. et al. Co-expression of CD39 and CD103 identifies tumor-reactive CD8 T cells in human solid tumors. Nat Commun 9, (2018).

17. Losurdo, A., et al. Single-cell profiling defines the prognostic benefit of CD39high tissue resident memory CD8+ T cells in luminal-like breast cancer. Commun Biol 4, (2021).

18. Bagaev, A. et al. Conserved pan-cancer microenvironment subtypes predict response to immunotherapy. Cancer Cell 39, (2021).

19. Eiva, M. A., Omran, D. K., Chacon, J. A. & Powell, D. J. Systematic analysis of CD39, CD103, CD137, and PD-1 as biomarkers for naturally occurring tumor antigen-specific TILs. Eur J Immunol 52, (2022).

20. Sun, W. et al. CD57-positive CD8 + T cells define the response to anti-programmed cell death protein-1 immunotherapy in patients with advanced non-small cell lung cancer. NPJ Precis Oncol 8, (2024).

21. Huang, B. et al. CD8 + CD57 + T cells exhibit distinct features in human non-small cell lung cancer. J Immunother Cancer 8, (2020).

22. Litwin, T. R. et al. Infiltrating T-cell markers in cervical carcinogenesis: a systematic review and meta-analysis. Br J Cancer 124, (2021).

23. Liu, P., Zhao, L., Kroemer, G. & Kepp, O. Conventional type 1 dendritic cells (cDC1) in cancer immunity. Biology Direct vol. 18 Preprint at 10.1186/s13062-023-00430-5 (2023).

24. Sisirak, V. et al. Impaired IFN-α production by plasmacytoid dendritic cells favors regulatory T-cell expansion that may contribute to breast cancer progression. Cancer Res 72, (2012).

25. Zhou, B., Lawrence, T. & Liang, Y. The Role of Plasmacytoid Dendritic Cells in Cancers. Frontiers in Immunology vol. 12 Preprint at 10.3389/fimmu.2021.749190 (2021).

26. Biswas, S. K. & Mantovani, A. Macrophage plasticity and interaction with lymphocyte subsets: Cancer as a paradigm. Nature Immunology vol. 11 Preprint at 10.1038/ni.1937 (2010).

27. Hristodorov, D. et al. Targeting CD64 mediates elimination of M1 but not M2 macrophages in vitro and in cutaneous inflammation in mice and patient biopsies. MAbs 7, (2015).

28. McGovern, N. et al. Human dermal CD14+ cells are a transient population of monocyte-derived macrophages. Immunity 41, (2014).

29. Prat, M. et al. Circulating CD14 high CD16 low intermediate blood monocytes as a biomarker of ascites immune status and ovarian cancer progression. J Immunother Cancer 8, (2020).

30. Rocher, B. Du, Mencalha, A. L., Gomes, B. E. & Abdelhay, E. Mesenchymal stromal cells impair the differentiation of CD14++ CD16- CD64+ classical monocytes into CD14++ CD16+ CD64++ activate monocytes. Cytotherapy 14, (2012).

31. Peng, Q. et al. PD-L1 on dendritic cells attenuates T cell activation and regulates response to immune checkpoint blockade. Nat Commun 11, (2020).

32. Meng, L. et al. Mechanisms of immune checkpoint inhibitors: insights into the regulation of circular RNAS involved in cancer hallmarks. Cell Death and Disease vol. 15 Preprint at 10.1038/s41419-023-06389-5 (2024).

33. Coombes, J. L. et al. A functionally specialized population of mucosal CD103+ DCs induces Foxp3+ regulatory T cells via a TGF-β -and retinoic acid-dependent mechanism. Journal of Experimental Medicine 204, (2007).

34. Iliev, I. D. et al. Human intestinal epithelial cells promote the differentiation of tolerogenic dendritic cells. Gut 58, (2009).

35. Togashi, Y., Shitara, K. & Nishikawa, H. Regulatory T cells in cancer immunosuppression — implications for anticancer therapy. Nature Reviews Clinical Oncology vol. 16 Preprint at 10.1038/s41571-019-0175-7 (2019).

36. Zhang, J. et al. The prognostic value of Th17/Treg cell in cervical cancer: a systematic review and meta-analysis. Front Oncol 14, (2024).

37. Jacobelli, S. et al. Anti-HPV16 E2 protein T-cell responses and viral control in women with usual vulvar intraepithelial neoplasia and their healthy partners. PLoS One 7, (2012).

38. La Gruta, N. L., Gras, S., Daley, S. R., Thomas, P. G. & Rossjohn, J. Understanding the drivers of MHC restriction of T cell receptors. Nature Reviews Immunology vol. 18 Preprint at 10.1038/s41577-018-0007-5 (2018).

39. Cai, H., et al. HPV16 E6-specific T cell response and HLA-A alleles are related to the prognosis of patients with cervical cancer. Infect Agent Cancer 16, (2021).

40. Krishna, S. et al. Human papilloma virus specific immunogenicity and dysfunction of CD8+ T cells in head and neck cancer. Cancer Res 78, (2018).

41. Halle, M. K. et al. A gene signature identifying CIN3 regression and cervical cancer survival. Cancers (Basel) 13, (2021).

42. Limones-Gonzalez, J. et al. Changes in the molecular nodes of the Notch and NRF2 pathways in cervical cancer tissues from the precursor stages to invasive carcinoma. Oncol Lett 28, 522 (2024).

43. Boutilier, A. J. & Elsawa, S. F. Macrophage polarization states in the tumor microenvironment. International Journal of Molecular Sciences vol. 22 Preprint at 10.3390/ijms22136995 (2021).

44. Haist, M., Stege, H., Grabbe, S. & Bros, M. Review the functional crosstalk between myeloid-derived suppressor cells and regulatory t cells within the immunosuppressive tumor microenvironment. Cancers vol. 13 Preprint at 10.3390/cancers13020210 (2021).

45. Salvo, G., Odetto, D., Pareja, R., Frumovitz, M. & Ramirez, P. T. Revised 2018 International Federation of Gynecology and Obstetrics (FIGO) cervical cancer staging: A review of gaps and questions that remain. International Journal of Gynecological Cancer vol. 30 Preprint at 10.1136/ijgc-2020-001257 (2020).

46. Jurtz, V. et al. NetMHCpan-4.0: Improved Peptide–MHC Class I Interaction Predictions Integrating Eluted Ligand and Peptide Binding Affinity Data. The Journal of Immunology 199, (2017).

47. Saini, S. K. et al. Not all empty MHC class I molecules are molten globules: Tryptophan fluorescence reveals a two-step mechanism of thermal denaturation. Mol Immunol 54, (2013).

48. Bentzen, A. K. et al. T cell receptor fingerprinting enables in-depth characterization of the interactions governing recognition of peptide–MHC complexes. Nat Biotechnol 36, (2018).

